# Zero-Shot Generation of Protein Conformational Ensembles Through AlphaFold Latent Flooding

**DOI:** 10.64898/2026.04.16.718914

**Authors:** Runtong Qian, Rui Zhan, Zilin Song, Jing Huang

**Affiliations:** State Key Laboratory of Gene Expression, School of Life Sciences, Westlake University, Hangzhou, Zhejiang 310030, China; Westlake AI Therapeutics Lab, Westlake Laboratory of Life Sciences and Biomedicine, Hangzhou, Zhejiang 310024, China

## Abstract

Despite transformative advances in protein structure prediction, generating conformational ensembles directly from sequence in an efficient and accessible manner remains a central challenge. We introduce a heuristic importance sampling framework for the zero-shot prediction of functionally relevant, conformationally diverse structures, bypassing the need for extensive physics-based modeling or prior domain knowledge. Guided by an analysis of massive activations in AlphaFold (AF), we develop a latent flooding algorithm that enables adversarial exploration of the latent space while preserving local geometric integrity. We show that the AF latent flooding (AFLF) method robustly recovers experimental structural fluctuations, captures function-relevant conformational states, and reveals cryptic binding sites across diverse protein systems. These results suggest that the AF latent space encodes functionally relevant conformational variability in a structured form that is readily amenable to *post hoc* exploration. Finally, we show that AFLF is computationally accessible and interoperable, offering a test-time generalization of AF for investigating protein dynamics and accelerating structure-based ligand discovery.

## 1 Introduction

Deep learning has catalyzed seminal advances in computational structural biology by facilitating accurate protein structure predictions and capturing essential biomolecular interactions. Exemplified by the AlphaFold (AF) series of models and related methods,^**1–6**^ these breakthroughs have delivered the predictions of static protein structures and their complexes with unprecedented accuracy.^**7**^ However, proteins are inherently dynamic entities that often adopt multiple conformational states in response to environmental perturbations or molecular interactions, a crucial aspect of their biological function.^**8**^

While methods based on neural networks have proven highly effective in predicting static, folded protein structures, they often struggle to enumerate the complete spectrum of functionally relevant states.^**9**^ This limitation significantly hinders the applicability of structural predictors like AF for crucial downstream tasks such as mechanistic investigation and allosteric ligand discovery.^**10**^ Recent advances have sought to bridge this gap by incorporating large-scale molecular dynamics (MD) simulations or leveraging deep generative methods to explore hidden protein conformations.^**11–13**^ Yet, these emerging approaches in general require substantial computational resources to generate labeled data and train models. The acquired capability is frozen in the released model, and extension to new purposes or interrogation of its inference mechanisms demands access to large-scale training data and hardware infrastructure that are not universally available. The nontrivial challenge remains the development of methods that can reliably generate diverse, biologically relevant structural ensembles solely from the primary sequence, while preserving the predictive accuracy and computational accessibility that have made biological foundation models like AF transformative.^**14**^

Feature perturbation strategies, including subsampling or clustering the multi-sequence alignments (MSAs), template manipulations, and test-time interventions, have nudged the AF models into producing alternative, biologically plausible conformations without retraining.^**15–24**^ Although approaches as such are demonstrably capable of expanding the conformational landscape accessible to a single sequence input, the stacked neural transformations render the underlying processes opaque to human perception. Consequently, the absence of mechanistic explanations for why and how such perturbation steers the network to predict new functional states precludes the conversion of AF latent space into an enumerable conformational repository, and its learned structural landscape is left beyond systematic exploration.

Here, we introduce AF Latent Flooding (AFLF), a heuristic multiscale framework that repurposes the AF2 model as a zero-shot engine for generating protein conformational ensembles. In contrast to conditioned prediction frameworks which steer the model outputs by user-defined restraints on geometric variables,^**21,22**^ AFLF explores the latent space through adaptive repulsion from previously visited conformations. The generated ensemble constitutes a set of candidate conformations rather than samples from a defined thermodynamic distribution, and such candidates can be directly exploited in downstream structure-based analyses. We dissect how the perturbations on the AF2 learned latent representations propagate into diverse structural outputs. We assemble key AF2 components to enable direct latent inference and integrate Low-Rank Adaptation (LoRA) for efficient protein structural enumeration. We develop a self-repelling, self-adaptive sampler to efficiently explore the broad yet relevant conformational space embedded in the AF2 latent landscape. Through applications including quantification of structural fluctuations, prediction of large-scale conformational changes, and detection of cryptic pockets, we demonstrate that AFLF enables highly accessible ensemble generation, extending the AF foundation models into readily deployable assets for structure-based ligand discovery and related downstream applications.

## 2 Results

### 2.1 Massive activations in AF latent representations

To probe the latent space in the AF2 model, we modified its inference pipeline to retrieve the latent features: the MSA/pair tensors fed to the Evoformer stack and the single/pair tensors fed to the structural module (StructMod). Interestingly, all four latent tensors exhibited the same “massive activation” pattern reported in transformer-based large language models (LLMs),^**25**^ where a minute fraction of tensor elements deviates by orders of magnitude from the median (**Figure 1a**). Empirically, the massively activated elements in general encode input-agnostic attention biases that dominate layer-wise attention probabilities, yet their magnitudes are strictly required for model performance. Recent works have shown that protein language models (PLMs) and LLMs adopt similar patterns of geometry in their latent representations.^**26,27**^ We therefore expect that the massive activations in AF2 serve an analogous function, as the Evoformer stack, which is trained with a masked language (MSA) modeling objective, effectively renders the model also a PLM.

**Figure 1.**
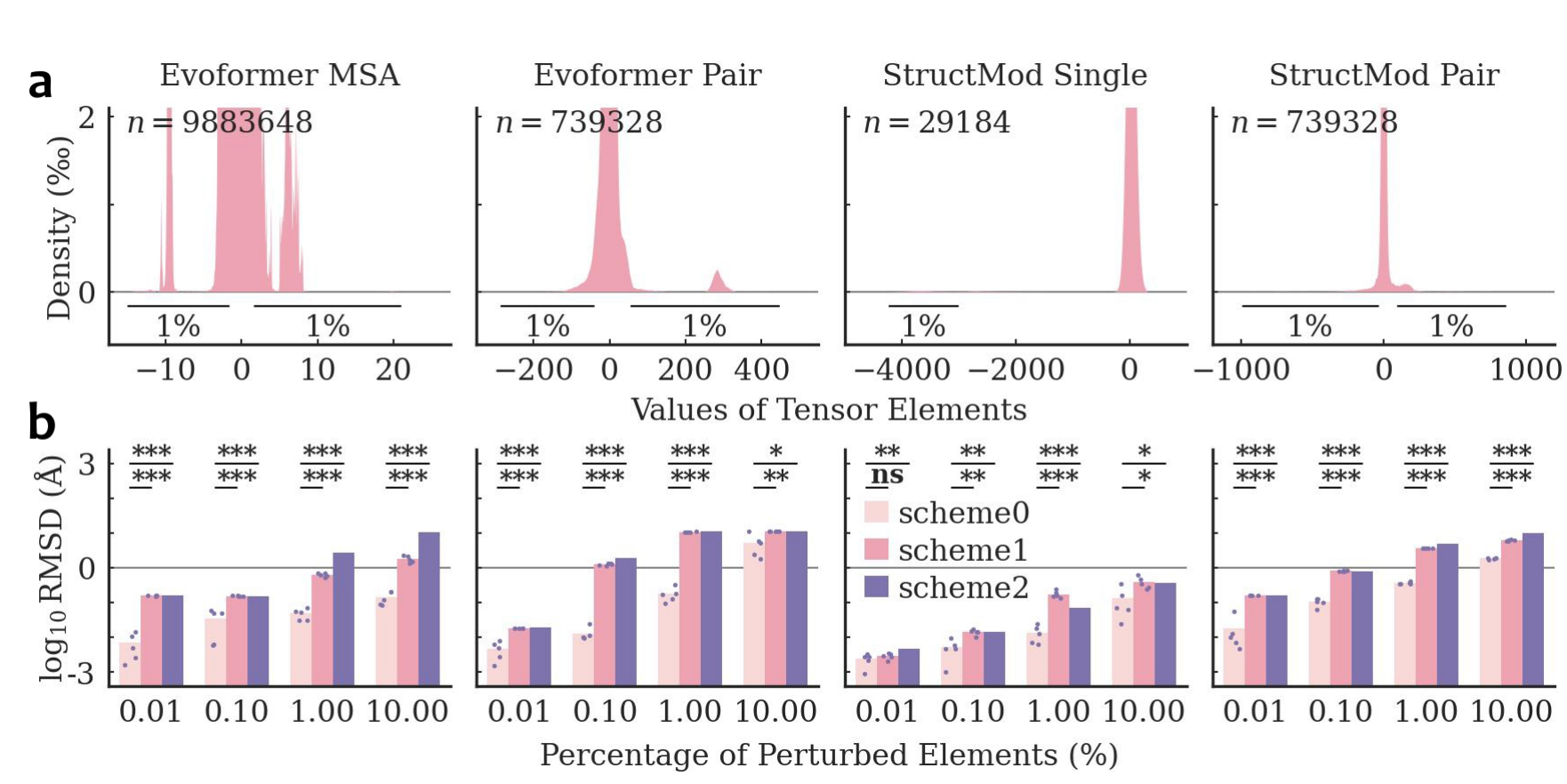
Latent features distribution and ubiquitin structural deviations from perturbed latent features. **(a)** Permillage value distributions of the AF2 latent tensor elements used to predict ubiquitin conformations. The 1st and 99th percentiles are indicated and *n* is the total number of elements in each tensor. **(b)** Backbone root mean squared deviations (log_10_RMSDs) of ubiquitin conformations predicted from perturbed latent tensors relative to the native unperturbed prediction. Statistical tests: scheme0 (*n* = 5) versus scheme1 (*n* = 5) by independent samples *t*-test; scheme0 (*n* = 5) versus scheme2 (*n* = 1) by one-sample *t*-test. **\*\*\***: *p <* 0.001; **\*\***: *p <* 0.01; **\***: *p <* 0.05; **ns**: not significant.

To delineate the contribution of the massively activated elements, we systematically ablated the four latent tensors. We flagged the tensor elements whose values fell within the extreme tails (top and bottom fractions) of the overall distribution as massively activated and subjected them to three perturbation schemes. Briefly, scheme1 resampled the flagged entries from a Gaussian distribution fitted to the full-tensor statistics and scheme2 zero-masked the same flagged entries. The reference scheme0 masked an equal number of randomly chosen entries to provide a baseline for the information loss. Across all four latent tensors and repeated random seeds, the targeted perturbations of the massive activations (scheme1 and scheme2) produced statistically significant structural deviations from the reference (scheme0, **Figure 1b**). A recent study on a community reproduction of AF2^**28**^ showed that global zero-masking of the pair representations severely depresses the template-modeling score on the predicted structures.^**29**^ As the geometric learning objectives of AF2 were imposed almost exclusively on the Structural Module, it is intuitive that its pair representations primarily encode the latent manifold of structural topologies. Here, we further show that corrupting only the massive pair activations, either in the Evoformer or in the StructMod, was sufficient to cause mode-collapse in the predicted coordinates, whereas distributing the same number of zeros at random preserved topologically plausible predictions (**Supplementary Figures S1, S2, S3**). In contrast, perturbing the massive Evoformer MSA activations warped the global fold and softened the local secondary structures, without precipitating structural collapse, and the reference scheme0 that randomly masked identical numbers of tensors elements maintained the overall fold architecture. Moreover, perturbations on the StructMod single representation, regardless of schemes, produced no discernible difference in the predicted structure. A similar mode of action is also observed for larger proteins (**Supplementary Figures S4, S5, S6, S7**).

Collectively, the latent tensors in AF2, like those of LLMs, contain a small subset of massively activated elements. Across the four latent representations, the massively activated entries serve distinct functions in the structural inference pipeline: corrupting them in the two pair tensors collapses the predicted coordinates; in the Evoformer MSA tensor it reshapes the global fold while preserving local topology; and in the StructMod single tensor it produces no measurable difference. Our perturbation experiments suggest that AF2 embeds the mechanism of determining specific folding patterns within the Evoformer MSA massive activations. In parallel, the null effect of perturbing the StructMod single representations indicates that the Evoformer stack progressively distils coevolutionary information from the MSA representation into the pair representation, such that, once the distillation is complete, the residual first-row MSA tensor (the single representation) becomes too uninformative to impact the structural output.

### 2.2 Latent Flooding for Protein Conformation Generation

To access the embedded protein structural state in the AF2 latent space, based on the above analysis, numerical interventions should preferably target the Evoformer MSA representations whose extreme valued elements constitute the decisive mechanism of folding. Our preliminary experiments indicated that the latent tensors cannot be universally partitioned into the massively activated and the residual subsets as the layered nonlinear mixing couples their contributions; therefore, the entire Evoformer MSA tensor was adapted to ensure system-agnostic behavior. The AFLF modeling proceeds in two consecutive stages: (1) AFLF checkpoint modeling, in which the AF2 embedding module fuses the MSA and the template input features into the latent MSA and pair representations that are cached via a modified AF2 inference pipeline; and (2) AFLF inference modeling, where we initialize the Low-Rank Adaptation (LoRA)^**30**^ tensors on the cached checkpoints and propagate the augmented representations through the Evoformer and the StructMod stacks to generate new structures (**Figure 2a**). We note that the identical latent inference mechanism can also be applied to any intermediate layer as the checkpoint.

**Figure 2.**
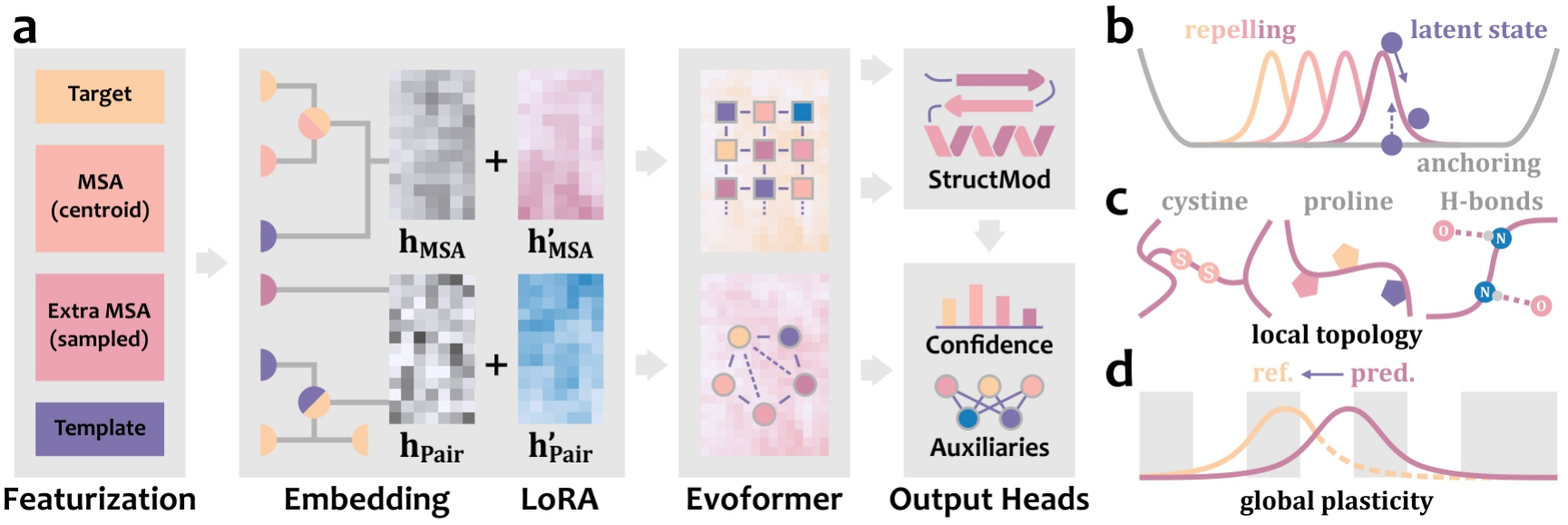
AlphaFold Latent Flooding (AFLF) model. **(a)** The AFLF model. **(b)** The repelling and the anchoring losses. The repelling loss drives the latent state away from previously visited conformations. The anchoring loss penalizes excessive deviation of repelled coordinates from their reference distances. **(c)** The local geometric loss enforces rigid-body restraints on conserved motifs via optimally aligned RMSDs. **(d)** The global geometric loss regulates protein-scale plasticity by minimizing the cross-entropy between reference and predicted inter-residue distance distributions in kernel space.

To explore the protein conformational landscape latent in AF, the AFLF inference modeling implements a self-repelling, self-adaptive enhanced sampling protocol (see **Methods**). The AFLF method characterizes the protein conformation by the inter-centroid distances between predefined atom groups, thereby providing a conformational space of reduced dimensions while retaining essential geometric relations. The AFLF sampler adopts a memory-guided strategy that maintains a running stack of previously visited repellent states which derives a series of Gaussian repelling potentials to penalize revisiting previous conformations (**Figure 2b**). To overcome the non-ergodicity inherent to naïve self-repelling walks in the AF latent space, AFLF implements an adaptive importance sampling scheme based on the coefficient of variations tracked along the trajectory. The adaptive coefficients scale inversely with the variation statistics: distant pairs exhibiting low variability are deemed insufficiently sampled and are assigned higher repulsive weights, whereas high variability distances with adequate sampling are downscaled. This self-adaptive mechanism dynamically redirects online the sampling effort toward underexplored regions of the latent conformational space.

To constrain the latent exploration on the manifold of native-like folds, AFLF incorporates a multiscale geometric regularization strategy comprising three complementary loss terms. The anchoring loss applies boundary penalties to prevent excessive deviations of repelled coordinates from their reference (**Figure 2b**). The local geometric loss enforces the structural integrity on essential motifs (such as disulfide bonds and proline rings) via optimally aligned root mean squared deviations (RMSDs, **Figure 2c**). The global geometric loss controls the protein-scale structural plasticity through the cross-entropy between the reference (defaulting to the AF naive prediction) and predicted inter-residue distance distributions (**Figure 2d**).

Conceptually, the online modulation of AFLF dynamics by the sampling statistics parallels the conformational flooding^**31**^ approach, in which Gaussian biases are planted along slow collective coordinate variations to accelerate classical MD sampling. The flooding bias in enhanced sampling MD acts on a physical potential energy surface, so the unbiased distribution can, in principle, be recovered through reweighting. The latent space of AF, by contrast, has no corresponding physical energy function or defined Boltzmann measure, and the repelling term provides the explicit driving force to navigate latent space in a manner that encourages conformational exploration. The AFLF continuously rescales the repulsive potentials according to the instantaneous sampling heterogeneity of the inter-centroid distances, thereby implementing a memory-guided adaptive enumeration of the AF latent manifold. The multiscale geometric regularization schemes (anchoring, local motif rigidity, and global distance distribution entropy) are fused with the adaptive repelling potential into a single differentiable objective. The gradient flow is realized by low-rank adaptation of the Evoformer MSA checkpoint, enabling direct exploration of the latent space without retraining the full network. AFLF optimization employs the same loss terms and the flooding potentials are applied at the structural scale that matches the motion of interest, residue-wise for local fluctuations and segment-wise for domain motions and pocket openings. Equipped with the AFLF protocol, we next apply it to reproduce experimental structural fluctuations, to explore alternative functional states, and to identify cryptic binding sites.

### 2.3 Generating Protein Structural Fluctuations

To validate AFLF for reproducing experimental structural fluctuations, we selected ubiquitin, a model protein whose high-resolution crystal structure exhibits a well-characterized flexibility gradient from a rigid core to a highly mobile C-terminus. The AFLF sampling was initialized from the latent state produced by the native AF prediction and was advanced for 20,000 steps with Gaussian flooding applied on all intercentroid distances between residues separated by at least four sequential positions, yielding a continuous conformational trajectory (**Supplementary Movie S1**). For both the MD baseline and the AFLF trajectory, we computed per-residue *B*-factors based on root mean square fluctuations (RMSFs).^**32**^

Remarkably, the structural flexibility of ubiquitin captured by AFLF reproduced the crystallographic *B*-factors at the same amplitude and preserved the ranking correlation as faithfully as the conventional MD reference: Kendall *τ* = 0.46 (~73% concordant pairs) for AFLF versus *τ* = 0.49 (~74.5% concordant pairs) for MD (**Figure 3a, Supplementary Table S1**). Visual inspection of the residue-level flexibility maps indicated that the AFLF ensemble reconstructed the experimentally observed pattern of rigid (purple) and mobile (pink) regions (**Figures 3b, 3c**). We also compute the ranking statistics excluding the flexible C-terminal tail (residues 72–76), which gives Kendall *τ* = 0.44 (~72% concordant pairs). In addition, we compared the AFLF ensemble with the experimental backbone N–H order parameters (*S*^2^) of ubiquitin from NMR relaxation,^**33**^ with the N–H vectors of the generated conformations reconstructed from the backbone heavy atoms, and found qualitative agreement in the overall profile (**Supplementary Figure S8**). We also extended the *B*-factor computation to nine additional proteins and benchmarked AFLF against BioEmu^**13**^ and AFsample2^**17**^ on this set (**Figure 3d, Supplementary Figures S9, S10, S11, S12, S13, S14, S15, S16, S17**). For all three methods, we computed the *B*-factors over four independent 5,000-frame trajectories. Notably, the AFLF values are the closest to the experimental magnitudes for the majority of the proteins, falling within a factor of two of the experimental values in eight of the nine cases. The two baselines deviate from experiment in opposite patterns: the *B*-factors from AFsample2 are systematically lower, whereas those from BioEmu are consistently higher, in the most pronounced cases by two orders of magnitude. In terms of ranking correlation, AFLF performs on par with BioEmu, a generative model trained on large-scale MD datasets, and exceeds AFsample2 on average (**Supplementary Figures S18**). Importantly, this agreement emerges without reweighting or recalibration, and is reached by AFLF in a zero-shot setting without any additional training.

**Figure 3.**
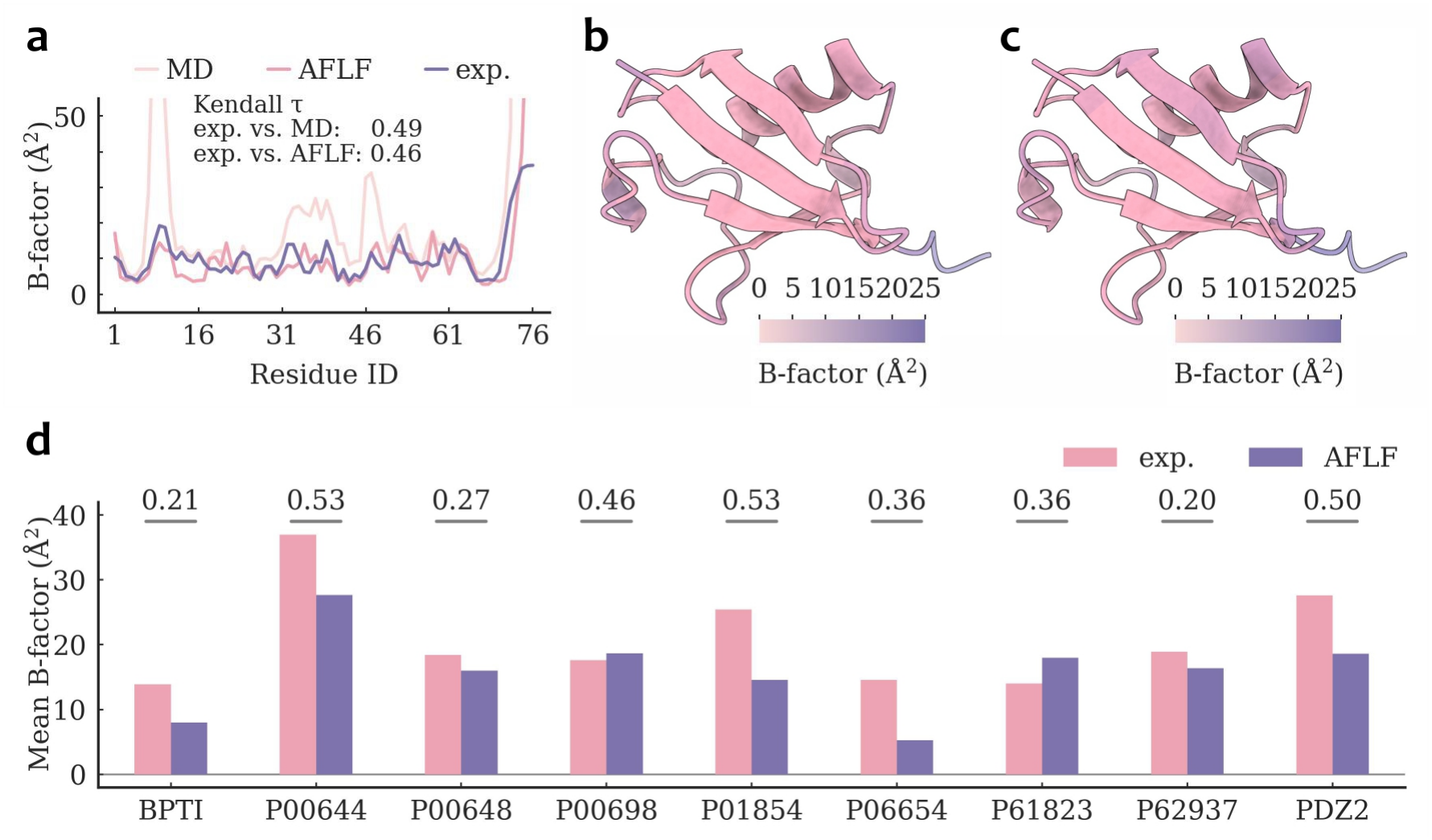
Structural fluctuations generated by AFLF. **(a)** The ubiquitin *B*-factors from the crystal structure (PDB id: 1UBQ), the baseline MD simulation, and the AFLF trajectory. The **(b)** AFLF-derived and **(c)** experimental ubiquitin structural fluctuations colored by low (pink) and high (purple) *B*-factors. **(d)** Mean *B*-factors of the selected proteins from experiment (pink) and from AFLF (purple), with the Kendall *τ* coefficients indicated above the bars. See also **Supplementary Figures S9, S10, S11, S12, S13, S14, S15, S16, S17, S18**.

### 2.4 Generating Protein Functional States

To evaluate AFLF for exploring large-scale conformational transitions, we focused on adenylate kinase (AdK), a model enzyme with well-characterized open and closed states, whose catalytic cycle involves the opening of the ATP-binding domain (LID, residues 121–159) and the nucleotide-monophosphate-binding domain (NMP, residues 30–59) with respect to the structurally rigid CORE domain (remaining residues).^**34**^ Beyond these two states, a locally unfolded state has been characterized experimentally as part of the native-state ensemble,^**35**^ although no atomic structure has been deposited for direct structural comparison. Starting from the AF-predicted closed conformation, we initiated a 10,000-step AFLF simulation after decomposing the protein into segments defined by the secondary structure elements: each *α*-helix, *β*-strand, and loop longer than six residues was treated as an independent segment. The AFLF flooding potentials were applied exclusively on the segment pairs whose minimal inter-residue distance fell below 4.5 Å and the non-contacting distances were left unbiased. The AFLF anchoring thresholds were set to −1 to +25 Å for contacting segment pairs, except for intra-domain pairs and those involving the CORE domain, which were restrained within −1 to +1 Å. Referring to the crystallographic closed and open structures, we projected the AFLF trajectory onto the two-dimensional LID-CORE/NMP-CORE plane for quantifying the AdK closed ↔ open transitions.

Notably, the AFLF ensemble populated a contiguous density of conformations between the closed and open crystallographic structures (**Figure 4a, Supplementary Movie S2**). The representative snapshots recovered both crystal structures with the C*α* RMSDs being 1.18 Å (closed, **Figure 4b**) and 3.06 Å (open, **Figure 4c**). Along the simulated trajectory, we recorded nine closed ↔ open transition events whose LID-NMP centroid distance traversed the full experimental interval (**Figure 4d**). The resulting structural distribution confirmed that the AFLF ensemble autonomously captured both native endpoints and interpolated the intermediate transient space that connects them. We further evaluated AFLF on the 44 domain-motion systems from the BioEmu benchmark,^**13**^ which undergo large-scale rearrangements of entire domains. Using AF2 as the underlying predictor and requiring no training stage, AFLF recovers the alternative experimental state in 30 of the 44 systems, approaching the 36 systems recovered by BioEmu (**Figure 4e, Supplementary Figure S19**). Among the representative systems, calmodulin rearranges its two globular lobes between the compact and the extended states, and AFLF recovers both endpoints of this classical domain motion (**Figures 4f, 4g, 4h, 4i**).

**Figure 4.**
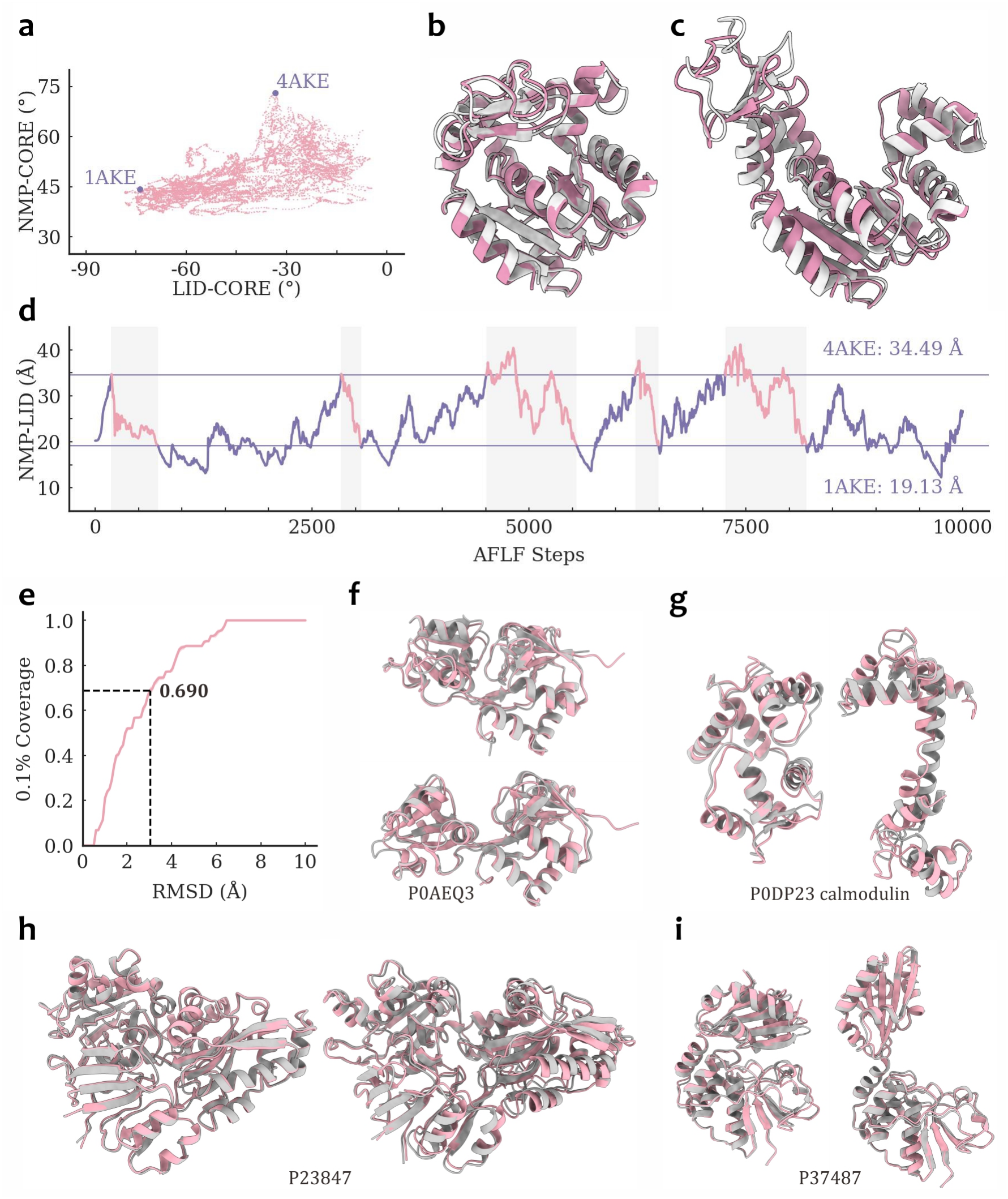
Functional states generated by AFLF. **(a)** The AFLF generated AdK trajectory projected on the two dimensional plane of the LID-CORE and NMP-CORE angles. The experimental closed (PDB id: 1AKE) and open (PDB id: 4AKE) structures are shown. The representative AFLF generated conformations (pink) for **(b)** the closed and **(c)** the open states are superposed with the experimental crystal structures (gray). **(d)** The AFLF generated AdK trajectory projected on the LID-NMP centroid distance. **(e)** The 0.1% coverage of the experimental alternative states on the 44 domain-motion systems from the BioEmu benchmark. The AFLF generated conformations (pink) that best match each of the two experimental states (gray) are shown for the representative systems **(f)** P0AEQ3, **(g)** P0DP23 (calmodulin), **(h)** P23847, and **(i)** P37487. See also **Supplementary Figure S19**.

### 2.5 Generating Protein Cryptic Cavities

To assess AFLF for detecting cryptic cavities, we targeted five proteins whose cryptic binding sites have been experimentally documented (**Supplementary Table S2**):^**36**^ the class A *β*-lactamase TEM-1, the anti-apoptotic protein Bcl-x_*L*_, the bovine *β*-lactoglobulin (*β*LG), the human nucleoside diphosphate kinase A (NDPK-A), and the human glycolipid transfer protein (GLTP). In all cases, the apo crystal structure captures the cryptic cavity in the occluded state and the exposed conformations emerge only under mutational or ligand-induced perturbations. A recent benchmark has shown that the native AF predictions capture both the occluded and the exposed cryptic binding site conformations only when the training data carries balanced populations of each,^**37**^ suggesting that the native AF inference pipeline is constrained to recapitulate the structural distributions already embedded in its learned repertoire. Here, we demonstrate that the AFLF approach effectively broadens the predicted conformational landscape and robustly predicts both the occluded and the exposed states without physics-based refinement. Similar to the AdK case, we began from the AF native predictions, partitioned the proteins into secondary structure segments, activated the AFLF flooding potentials for contacting segment pairs, and retained the unbiased treatment of non-contacting pairs. The anchoring bounds were uniformly set to −1 Å and +10 Å. Using the crystallographically resolved conformations as structural references, we monitored the volume of the cryptic cavities throughout the 10,000-step AFLF trajectories for all five test proteins.

Conspicuously, the AFLF ensembles spanned the full volumetric space between the apo-occluded and the holo-exposed conformations in all five test proteins (**Figure 5a**), indicating that the model did not merely recognize the two endpoint states but also captured the intervening transient states. In parallel, the C*α* RMSDs relative to both the apo and the holo states consistently remained below ~3 Å (**Figure 5b**), indicating that the sampled conformations preserved native folds. At the same time, the trajectories effectively accessed conformational states in which the cryptic pockets are fully exposed. Detailed inspection of the AFLF snapshots that best superimposed on the crystallographic conformations confirmed that the model recovered both states without being preconditioned on the ligand moiety (**Figures 5c, 5d, Supplementary Figure S20**). The minimal structural deviations (C*α* RMSDs *<*2 Å) and the close resemblance of the optimally aligned AFLF conformations to the holo states demonstrated that the method produced apo-exposed conformations that are immediately exploitable for structure-based ligand discovery.

**Figure 5.**
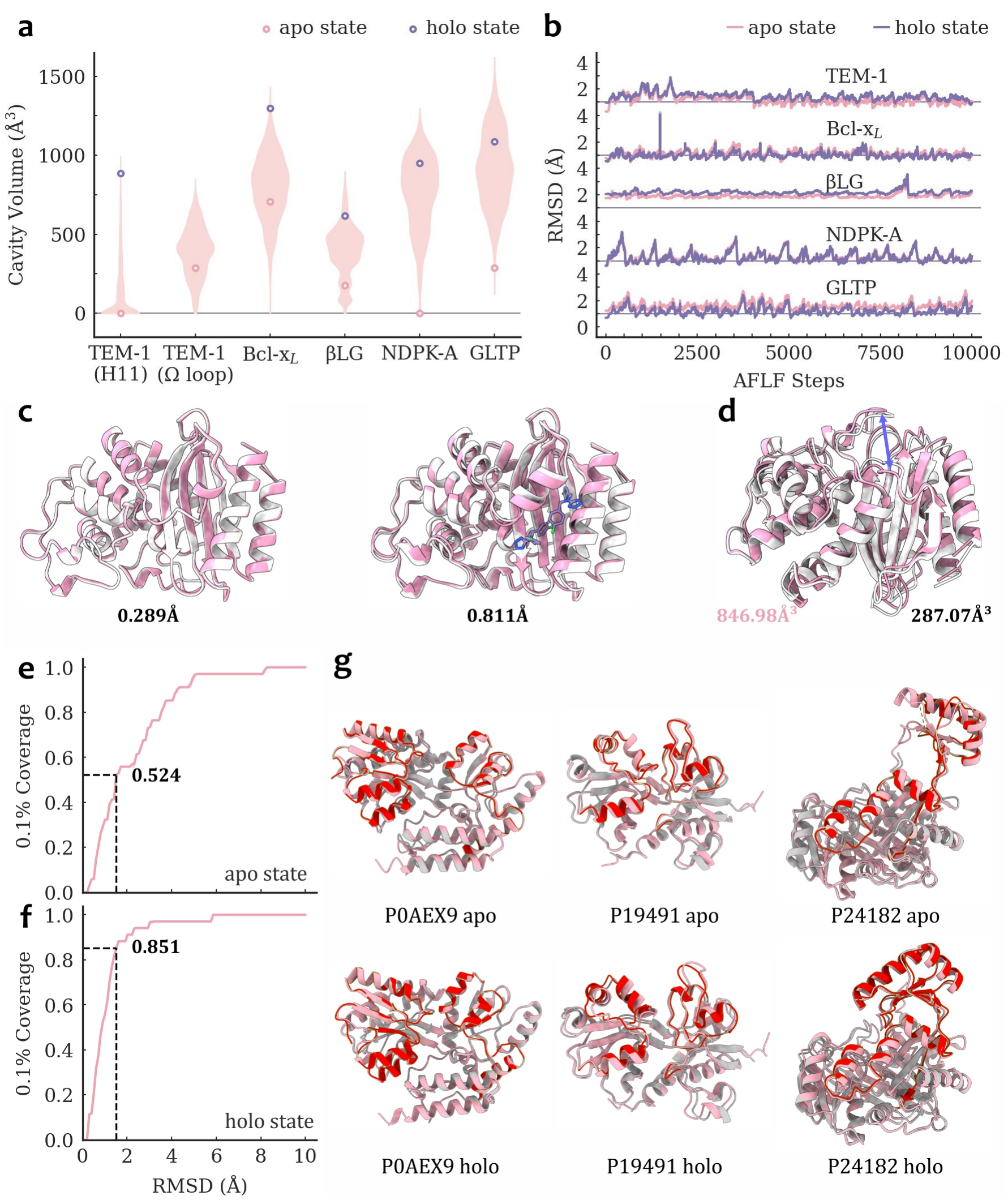
Cryptic cavities generated by AFLF. **(a)** The cavity volume distributions of the AFLF generated conformations. **(b)** The backbone C*α* RMSD profiles to the reference apo and holo conformations. The representative AFLF generated structures of the class A *β*-lactamase TEM-1 for **(c)** the apo state and the holo state that exposes the H11 cryptic binding site. The minimum backbone C*α* RMSDs between the AFLF generated conformations (pink) and the experimental crystal structures (proteins in gray and ligands in purple) are listed. **(d)** The AFLF generated TEM-1 conformation (pink) that exposes the Ω loop cryptic binding site (purple arrow), the apo state TEM-1 crystal structure (gray), and their cavity volumes. The 0.1% coverage of the experimental **(e)** apo and **(f)** holo states by the AFLF ensembles on the 34 cryptic-pocket systems from the BioEmu benchmark. **(g)** The AFLF generated conformations (pink) that best match the experimental apo and holo structures (gray) of the representative systems P0AEX9, P19491, and P24182, with the regions used by the BioEmu benchmark for structural coverage highlighted in red. See also **Supplementary Figures S20, S21, S22**.

Among the five systems, TEM-1 is an important therapeutic target for combating antimicrobial resistance, and its cryptic binding sites remain challenging to identify using either large-scale MD simulations or AF-based workflows^**37–39**^. TEM-1 has two reported cryptic binding sites. The H11 site, located between helices H11 and H12, has been structurally characterized in its exposed form (**Figure 5c**). In contrast, the site proximal to the Ω loop has not yet been experimentally observed in an exposed conformation, despite functional evidence supporting its role in allosteric regulation^**40**^. AFLF successfully identified both cryptic sites. For the H11 site, the majority of the sampled conformations did not contain this cavity, while full opening was captured as a rare event along the AFLF trajectory. For the Ω loop site, AFLF sampled an exposed state (**Figure 5d**), with the cavity volume increasing from 287.07 Å^**3**^ to 846.98 Å^**3**^. Together, these results underscored AFLF’s ability to extrapolate plausible functional conformations beyond the training data based on experimentally resolved structures. To quantify the coverage of the experimental states beyond these five proteins, we evaluated AFLF on the 34 cryptic-pocket systems from the BioEmu benchmark,^**13**^ in which the conformational change is scored by the local backbone RMSD over the curated cryptic-site residues. At the threshold of 1.5 Å defined by BioEmu, AFLF recovers the holo and the apo states with 0.1% coverages of 0.851 and 0.524, respectively, which is exemplified by the representative systems (**Figures 5e, 5f, 5g**). AFLF therefore closely matches the metrics computed from the released BioEmu ensembles (0.882 and 0.543) on this benchmark (**Supplementary Figures S21, S22**). Notably, both methods achieve higher coverage for the holo states than for the apo states, an asymmetry that likely originates from the ligand-bound conformations prevalent in the structural training data rather than from the generative mechanisms.^**37**^

## 3 Discussions

The unprecedented progress made in deep-learning protein structure prediction has culminated in biological foundation models that deliver coordinates at atomic resolution directly from the primary sequence. However, the algorithms of single-state predictors such as AF were optimized to reproduce the most probable conformations in the protein structural database and not the geometrical heterogeneity underlying catalysis, allostery, or ligandability. Here, we provide AFLF, a democratized and interoperable ensemble generation protocol that autonomously explores the structural landscapes encrypted within the AF latent feature space.

To dissect how the latent dynamics driven by AFLF evolves the massive activations, we traced every latent state visited along the AdK trajectory. For each AFLF step, we extracted the (top and bottom) 1% most extreme entries in the Evoformer MSA representation, computed the Jaccard index (JI) between each pair of latent states, and embedded the latent trajectory using the precomputed JI distance matrix with the Uniform Manifold Approximation and Projections (UMAP) method (**Figure 6a**). Although the AdK backbone repeatedly transitioned between the open and closed conformations, the latent trajectory followed a unidirectional path in the UMAP embedding, with no cyclic pattern observed (**Figures 6b, 6c, 6d, 6e**). A consistent monotonic trend was observed when the optimal-transport distance (*d*_OT_) between each latent state and the initial state was plotted against the AFLF step, with *d*_OT_ increasing almost linearly along the trajectory (**Figures 6a**,**Supplementary Figure S23**). Evidently, the identity and the magnitude of the massive activations continuously evolve throughout the AFLF dynamics whereas the structural output can revert to an earlier macroscopic state. The many-to-one mapping of the latent states to the structural outputs justifies our choice of operating LoRA on the entire MSA tensor rather than selectively on the initial massive activation set. In addition, the dynamical pattern also explains why earlier heuristic approaches – such as MSA subsampling or clustering – succeed only sporadically: these methods rely on manipulating the discrete MSA inputs and lack the continuously tunable latent coordinates to systematically traverse the AF latent manifold of topologically accessible ensemble of conformations. AFLF, by contrast, bypasses MSA curation altogether and re-tunes its perturbations in response to intermediate sampling feedback, providing a stable, on-the-fly enumeration through the latent conformational ensemble. A complementary strategy is pursued by the recent AF-RandomWalk, which diversifies the AF outputs by applying undirected stochastic noise to the network weights.^**23**^ Concurrent with our work, ConforNets demonstrated that the latent space can be explored with a single repelling-type objective optimized at inference time.^**24**^ In contrast, AFLF exploits the latent representation through a self-guided, self-regularized optimization, with the flooding potentials constructed according to the characteristics of the latent representations identified above.

**Figure 6.**
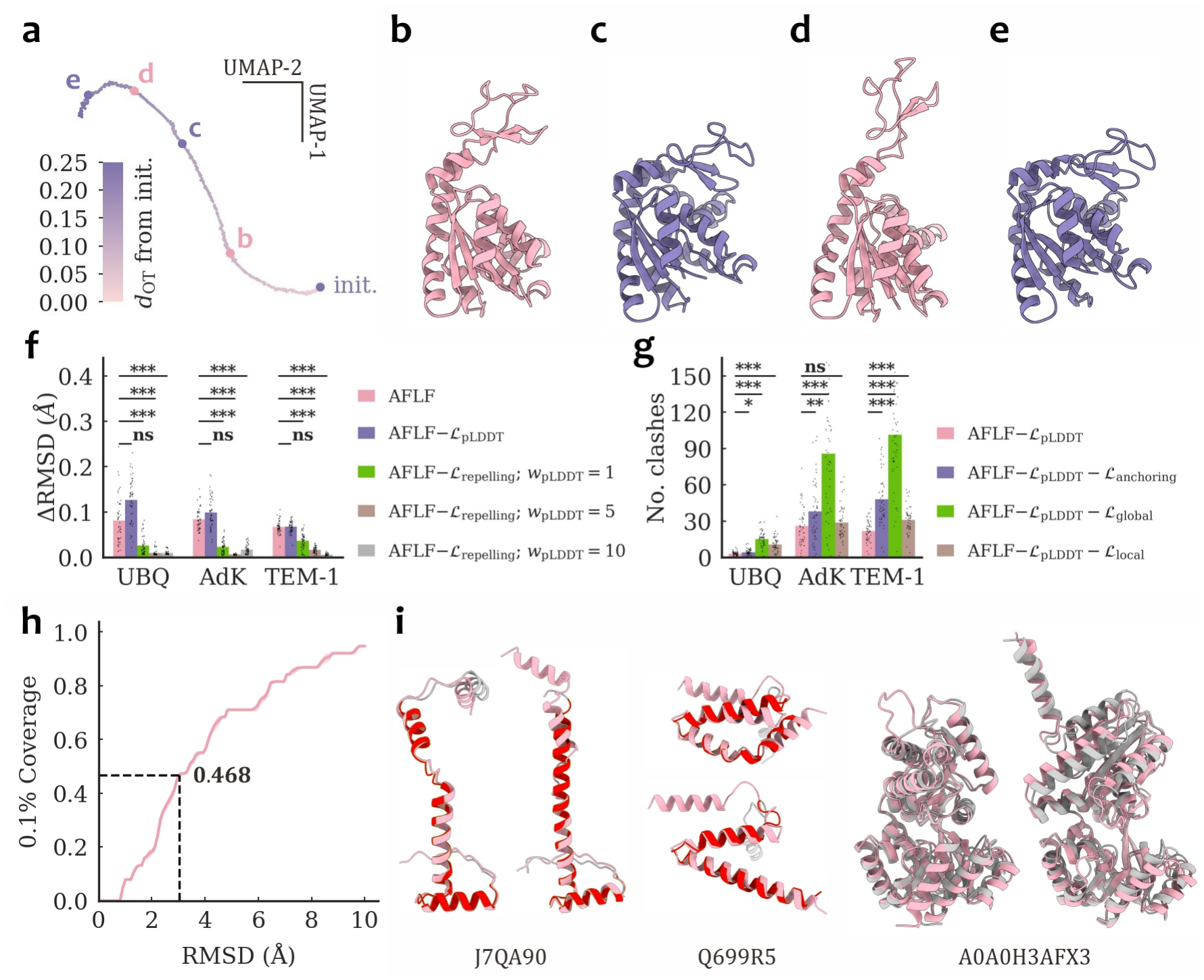
AFLF latent dynamics analysis, ablation experiments on representative systems, and out-of-distribution predictions. **(a)** The AFLF latent trajectory of AdK projected on the two-dimensional Uniform Manifold Approximation and Projection (UMAP) embedding using the Jaccard index as the distance metric. All latent states are colored by their optimal transport distances (*d*_OT_) to the initial state. **(b)**–**(e)** The AdK conformations generated from the four latent states marked in **(a)**, shown in the same colors as the corresponding latent states. **(f)** Per-frame backbone structural changes (ΔRMSD) along trajectories generated by the full AFLF objective and its ablated variants, including the no-repelling controls with the confidence term retained at *w*_pLDDT_ = 1, 5, and 10, for ubiquitin, AdK, and TEM-1. **(g)** Numbers of steric clashes in the conformations generated with the confidence term uniformly disabled, and with the anchoring, the global, or the local term additionally disabled, to expose the geometric effects of AFLF. In **(f)** and **(g)**, AFLF ℒ_X_ denotes the ablated variant with the loss term ℒ_X_ disabled. Each trajectory was partitioned into consecutive non-overlapping windows of 200 frames, yielding *n* = 50 window-averaged samples for each system. Bars indicate the mean across the window-averaged samples, and dots the individual values. Statistical tests compare each ablated variant against the full AFLF objective in **(f)** and against AFLF ℒ_pLDDT_ in **(g)** by independent samples *t*-test. **\*\*\***: *p <* 0.001; **\*\***: *p <* 0.01; **\***: *p <* 0.05; **ns**: not significant. **(h)** The 0.1% coverage of the target states on the 38 structures from the BioEmu OOD-60 benchmark. **(i)** The AFLF generated conformations (pink) superposed with the experimental target structures (gray) of three representative systems J7QA90, Q699R5, and A0A0H3AFX3, with the regions used by the BioEmu benchmark for structural coverage highlighted in red. See also **Supplementary Figures S23, S24, S25, S26, S27**.

Technically, four lines of evidence collectively establish the mechanistic validity and the empirical effectiveness of the AFLF protocol. First, the systematic ablation of the four AF latent tensors reveals that the massively activated elements in the Evoformer MSA representation act as the decisive fold maker, whereas corruption of the pair tensors precipitates coordinate collapse and perturbation of the StructMod single tensor is effectively silent. Second, the AFLF flooding potentials are modulated through the statistical variation rescaling to circumvent the non-ergodicity that stalls the naïve self-repelling walks and distances that have not yet varied are automatically penalized, ensuring that the sampler schedule allocates computational efforts to under-explored regions rather than oscillating within a local basin. Third, the interoperability of AFLF is substantiated by its consistent performance across spatial resolutions: residue-level inter-centroid distances recover crystallographic *B*-factors, whereas segment-level distances capture functional state transitions and expose cryptic binding sites. Fourth, an ablation study on representative systems clarifies the distinct roles of the AFLF loss terms. Removing the ℒ_repelling_ term abolishes conformational exploration entirely even with the ℒ_pLDDT_ term active, and increasing the confidence weight up to tenfold does not restore exploration, indicating that the confidence objective alone cannot drive the latent state away from its initial values (**Figure 6f, Supplementary Figures S24, S25**). Alternatively, omitting the anchoring, the local, or the global regularization term leads to structural distortions at the corresponding spatial scale, reflected in elevated steric clashes (**Figure 6g, Supplementary Figure S26**). The full objective thus constitutes the minimal combination required for stable latent-space sampling, while evaluation of these loss terms adds negligible computational overhead as the per-step cost is dominated by network inference.

To test whether AFLF recovers conformational states that AF2 did not observe during training, we evaluated AFLF on the OOD-60 set from the BioEmu benchmark,^**13**^ which comprises 38 systems whose reference structures were deposited in the PDB after the AF2 training cutoff and whose sequences share less than 60% similarity with any protein in the AF2 training set. On this held-out set of 38 systems, AFLF recovers the target states in 17 cases, compared with 23 systems recovered in the released BioEmu ensembles (**Figures 6h, 6i, Supplementary Figure S27**). Given this construction, and since the AFLF geometric losses encode no information about any particular target state, recovery of these states cannot be entirely attributed to memorization. Instead, these results indicates that the AFLF latent exploration can generalize beyond the structural instances its baseline AF2 was trained on and recover conformations that are not accessible using standard inference.

Moreover, the handful of transient cryptic cavities documented to date represent only a minute fraction of the proteome and remain numerically overwhelmed by the orthosteric sites that dominate contemporary structure-based ligand design. Here, AFLF produces conformational ensembles without recourse to seeded ligand poses, mutational profiles, or coevolutionary restraints; the protocol recovers the structural ensemble solely by proposing potential configurations that are intrinsically sustainable by the apo architecture. As the AFLF inference is decoupled from retrospective knowledge, every cavity it reports is a topological attribute distilled from the structural fluctuations embedded in the AF model rather than the artifact of inductive bias. We emphasize that the AFLF output is an unweighted set of candidate conformations, and the method does not assign Boltzmann weights to the generated structures. Such candidates are useful in settings where the primary bottleneck is obtaining physically reasonable conformations rather than determining their equilibrium populations. Examples demonstrated in this work include the recovery of functional states that can explain experimental observations and the exposure of cryptic binding sites that can be directly exploited for ligand discovery. When thermodynamic populations are required, selected AFLF conformations can further serve as starting points for physics-based sampling and free energy calculations. AFLF is complementary to specifically trained ensemble generators such as BioEmu, whose conformational knowledge is distilled from large-scale MD datasets through a dedicated training stage. AFLF instead decodes the structural variability already encoded in a pretrained predictor, so the general consistency of the two approaches on the majority of the benchmark sets above reflects independent lines of evidence. In addition, the AFLF sampling proceeds as a continuous trajectory that connects the recovered states through the intermediate conformations visited along the way, whereas trained ensemble generators or perturbative methods produce independent samples that do not guarantee such continuous evolution. AFLF involves no training stage and operates directly on a pretrained structure predictor, which makes it applicable when training resources or simulation data are unavailable, when a specific type of motion is of interest, or when newly released structure prediction models are to be used directly.

In summary, we present AFLF to bridge the gap between the single state predictions of AF and the conformational variability that underlies protein function. By grafting a lightweight, self-repelling, self-adaptive importance sampler onto the LoRA perturbations to the Evoformer latent representation, we transform the otherwise opaque AF latent embeddings into a navigable manifold and generate topologically accessible ensembles at a moderate computational cost. The AFLF protocol offers a fully modular in-place augmentation to AF, thereby delivering an immediately deployable enhancement to the foundational model without *post hoc* fine-tuning. Because no training stage is involved, the AFLF strategy can in principle be migrated to other structure prediction models, including those not designed or trained for ensemble generation, and can inherit improvements in the underlying model without additional training. Extending AFLF to more recent architectures such as AlphaFold3 would enable latent-space conformational sampling to RNA, whose dynamics are of great functional importance yet remain poorly characterized. Of course, combining physics-based modeling with AFLF may still confer measurable advantages in selected scenarios, such as determining the optimal transition paths.^**41**^ Finally, beyond its immediate practical utility, the AFLF protocol also furnishes a conceptual template for transforming a discriminative foundation model into a generative engine that continuously produces diverse conformational hypotheses. The same “latent flooding” philosophy can, in principle, be transposed to AF-Multimer to sample protein-protein complexes or to co-folding frameworks for exploring biomolecular interactions.

## 4 Methods

### 4.1 AFLF Checkpoint Modeling

To retrieve the initial latent states from AF, we execute a lightly modified AF inference pipeline and collect the latent input representations to the Evoformer, 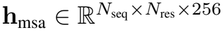 and 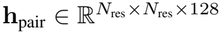, as well as to the StructMod, 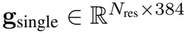 and 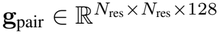. For each of the representations, we instantiate the LoRA perturbative tensors (rank = 8) from the uniform distribution (0.1, 0.1). The initially adapted latents are,

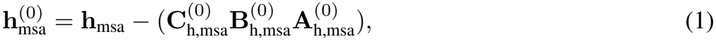

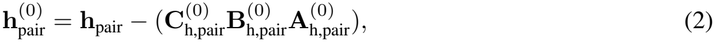

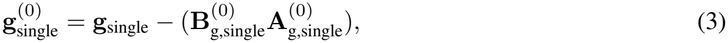

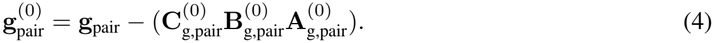

Using the LoRA tensors as the input, the AFLF inference pipeline can use the input checkpoint to the Evoformer or the StructMod as the forward entry. To infer from the Evoformer checkpoint, we use,

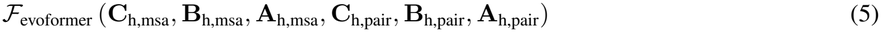

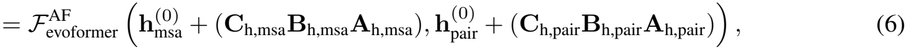

where 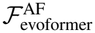 is the AF native inference function from the Evoformer inputs. To infer from the StructMod checkpoint, we use,

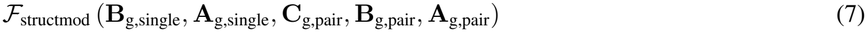

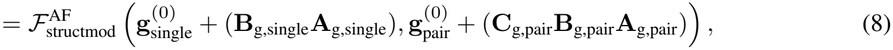

where 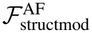 is the corresponding AF native inference function from the StructMod inputs.

### 4.2 AFLF Inference Modeling

In this study, without loss of generality, the AFLF model applies LoRA selectively to the Evoformer MSA representations,

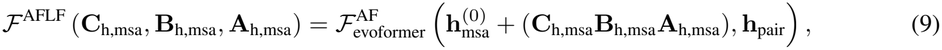

and outputs the coordinates **r** ∈ ℝ^*N×*3^ as well as the confidence scores 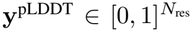. The output heavy atom coordinates from the main AF trunk are end-to-end differentiable with regard to all inputs, including the LoRA tensors introduced by AFLF. Accordingly, we operate most of the AFLF objectives on the pairwise distances between the centroids of the *M* atom groups formed by partitioning the total number of *N* predicted atomic coordinates. Practically, the user-defined lookup table *G* := {*G*_*i*_ | *i* ∈ {0, …, *M* − 1}} maps the heavy atom *n* ∈ *G*_*i*_ with coordinates **r**_*n*_ to the *i*-th atom group, for which the centroid coordinates **c** ∈ ℝ^*M×*3^ are computed,

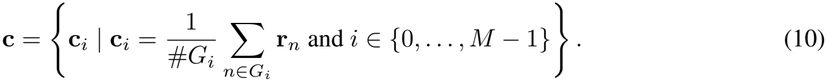

The AFLF structural state can then be characterized by the ordered set of inter-centroid distances,

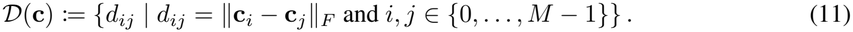

The notation is also used explicitly on the predicted atomic coordinates, *D*(**c**) = *D*(**r**; *G*) =: *D*_*G*_ (**r**).

To facilitate the AFLF structural exploration, we propose a self-repelling, self-adaptive walk guided by the Gaussian flooding potentials in the continuum of the AFLF inferred conformations. Let *τ* be the instant (artificial) time and Δ*t* the interval of taking repelling states, we keep a running stack of *K* memorized repellents {*D*_*G*_ (**r**(*t*_*k*_)) | *k* ∈ {0, …, *K* − 1}} taken at time stamps *t*_*k*_ = ⌊max(0, *τ/*Δ*t* − *k*)⌋ Δ*t*, where ⌊*·*⌋ denotes the floor rounding function. The flooding potential contributed by each of the repellent distances *d*_*ij*_(*t*_*k*_) ∈ *D*_*G*_ (**r**(*t*_*k*_)) on the instantaneous AFLF prediction *d*_*ij*_(*τ*) ∈ *D*_*G*_ (**r**(*τ*)) reads,

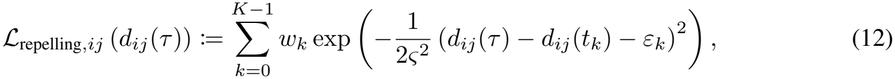

where *w*_*k*_ denotes the weight per repellent, *ς* the Gaussian width, and *ε*_*k*_ ~ *U* (−*ϵ, ϵ*) the perturbative noise capped at *ϵ* to prevent instantaneous (*τ* = *t*_*k*_) gradient vanishing. The reduced repelling loss is developed,

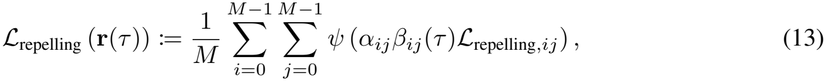

where *α*_*ij*_ ∈ {0, 1} is the predefined interaction mask, *β*_*ij*_(*τ*) ℝ_*>*0_ the self-adaptive coefficient, and *ψ* the random element-wise dropout function.

The self-adaptive coefficient *β*_*ij*_(*τ*) is of particular interest and deserves further discussion. Prior works have postulated that AF models internalize protein conformational variability from the protein structural data on which they were trained.^**42,43**^ However, the structural ensemble generated by mutating AF latent states captures neither realistic nor artificial times, and carries no thermodynamic weights. Consequently, the dynamics on the latent landscape of the AF foundational model lacks the ergodic freedom needed for basin escaping. Prolonged naïve self-repelling samplers therefore in general fail to reproducibly explore diverse protein conformations. While the AF models *per se* are not tied to physical temporal scales, we propose that the structural ensembles learned by AF can nevertheless be decoded by enhanced sampling across multiple spatial scales. The AFLF sampling is enhanced by a self-adaptive scheme facilitated through online importance rescaling, as detailed below. Taking the arithmetic average and the variance of each *d*_*ij*_ along the AFLF trajectory, 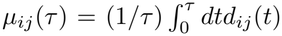and 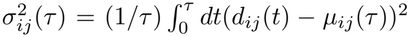, we estimate the instantaneous sampling progress using the *α*_*ij*_-masked coefficient of variation (CV),

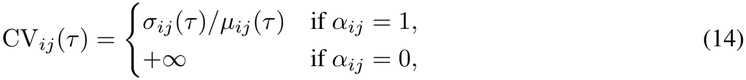

and define,

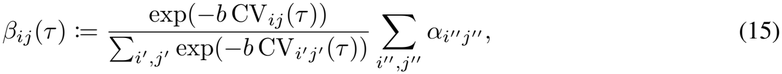

where *b* ∈ ℝ_⩾0_ is the reciprocal softmax tempering parameter and the constant scaling 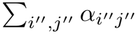 equates to 1 the average of all *β*_*ij*_(*τ*) that are unmasked (*α*_*ij*_ = 1). Briefly, distances with low CV_*ij*_(*τ*) are deemed under-sampled and are driven to explore further by a stronger Gaussian penalty with large *β*_*ij*_(*τ*), and *vice versa*. The tempering parameter *b* controls how peaked the softmax distribution is. For example, taking *b* → + ∞ would focus the sampling only on the lowest fluctuating *d*_*ij*_ whereas *b* = 0 leads to uniformly weighted sampling on all *d*_*ij*_. It should be noted that the *β*_*ij*_(*τ*)-modulated importance sampling implicitly assumes homogeneous thermal distribution among all *d*_*ij*_, ignoring that, for example, the distance between the centroids of two *α*-helices is intrinsically less variable than one spanning two disordered loops. In cases where the relevant {*d*_*ij*_} exhibit nontrivial heterogeneous scales in terms of their physical distribution, in addition to calibrating *β*_*ij*_(*τ*) for selected entries *a priori*, one could instead apply the centroid-wise rescaling,

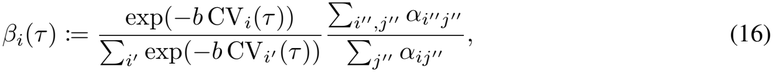

where 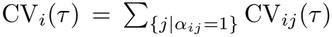 and ∀*j* : *β*_*ij*_ (*τ*) = *β*_*i*_ (*τ*). We note that adopting *β*_*i*_ (*τ*) entails that the centroids concerning fewer distances 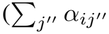 small) are permanently assigned proportionally larger rescaling weights. When {*α*_*ij*_} is derived from local interaction networks such as contact maps, the *β*_*i*_(*τ*) rescaling mirrors the empirical knowledge that higher contact density stabilizes local conformational dynamics. Of course, an unbiased measure of “sampling adequacy” — were it available without further unjustified statistical or thermodynamical hypothesis — would be the ideal metric for computing *β*_*ij*_(*τ*) online. In practice, the CV-derived estimators add negligible *O*(1) computational overhead and are likely sufficient for many applications. Relatedly, the current formulation operates on pairwise distances between individual residues or intact segments and is therefore not designed to capture partially folded or unfolded states involving disordered secondary structure shifts.

To contain the AFLF LoRA-ed latent states on the manifold of native-like folds, we introduce a multiscale strategy for regulating the structural outputs. We note that all of the following loss terms targeting distinct spatial scales can be selectively disabled according to the intended scope of application.

On the spatial scale of the atoms, the local structural features should be preserved by rigid body restraints for all AFLF generated conformations. Denote **r**_*p*_(*τ*) as the subset of atomic coordinates that sufficiently characterizes the *p*-th local spatial feature and 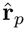 the corresponding reference from, e.g., the initial prediction **r**(0). The local topological loss is the optimally aligned deviation of the *P* subsets from the reference,

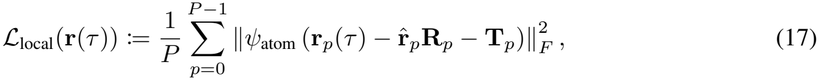

where *ψ*_atom_ denotes the atom-wise dropout and 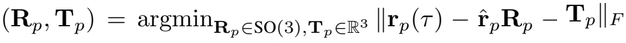. We compute each (**R**_*p*_, **T**_*p*_) pair with the closed-form Kabsch solver,^**44**^ which does not need to be differentiable since the optimal alignment acts only on the constant reference. Typically, we apply ℒ_local_ to maintain the structural constraints of molecular topologies that are not explicitly encoded by the internal topology used in AF inference, such as the disulfide bonds between cysteines, the pyrrolidine ring in prolines, and essential hydrogen bonding interactions.

On the spatial scale of the residues, the global structural plasticity could be controlled according to the AF predicted native fold. We represent each residue by the coordinates of its centroid and collect a total number of *Q* inter-centroid distances from the instantaneous prediction 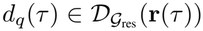 and the reference structure 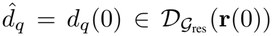, where _res_ is the residue-wise lookup table. The global plasticity loss is the cross-entropy between the density estimate on the reference and the prediction,

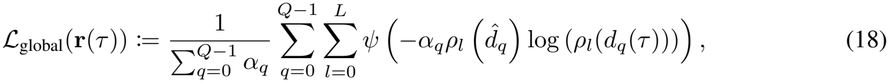

where *α*_*q*_ ∈ {0, 1} is the predefined binary mask that discards selected entries. Here, *ρ*_*l*_ is the kernel density estimator obtained from the histogram of *L* + 1 bins. By partitioning the support of the distance ensemble into bins of uniform width Δ*d* ∈ ℝ_*>*0_, we denote the *l*-th bin as [*l*Δ*d*, (*l* + 1)Δ*d*), except for the last bin [*L*Δ*d*, +*∞*). The distance density is estimated through the Gaussian kernel,

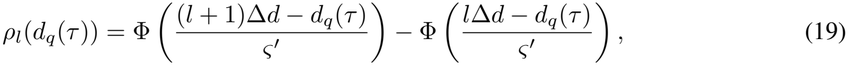

where 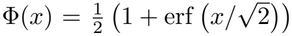 is the differentiable Gaussian cumulative density function and *ς*^*′*^ the smearing width. Typically, given arbitrary upper bound Δ*d*, we adapt *ς*^*′*^ to control the tolerated residue-wise fluctuations, and thus effectively the global structural plasticity. We note that ℒ_global_ is computed from kernel density estimations on calculated geometrical distances, as opposed to the softmax normalized latent representation in the “distogram” learning of AF.

On the spatial scale where the repelling potential is applied, it is possible that some of the repelling distances are not controlled by ℒ_local_ and ℒ_global_. We thus introduce the boundary penalty for each repelling distance to prevent large deviations 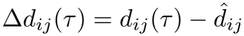 from the reference 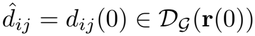,

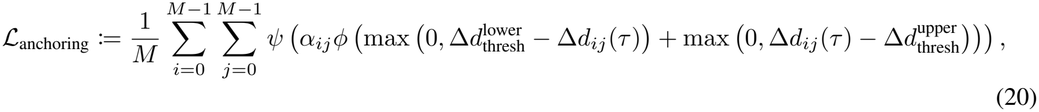

where *ϕ* denotes the Huber loss function^**45**^ and 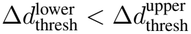 the lower and the upper bounds between which the boundary loss is inactive. Typically, ℒ_anchoring_ sets the permitted space of conformations for AFLF to explore. The anchoring bounds act as limits on the permitted structural deviation rather than target values toward which the optimization is driven. A bound set below the scale of the motion of interest prevents that motion from being sampled, whereas an overly permissive bound would reduce the sampling efficiency without altering the accessible conformations. The anchoring bounds can therefore be chosen from a coarse estimate of the motion magnitude, without knowledge of any target structure.

Finally, the AF model confidence loss ℒ_pLDDT_ := (1*/N*_res_) ∑**1** − **y**_pLDDT_ can be combined with the geometry losses into the time-dependent AFLF loss,

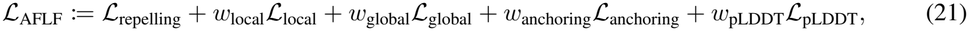

where *w*_local_, *w*_global_, *w*_anchoring_, and *w*_pLDDT_ are the corresponding scalar weights. Dynamical propagation on ℒ_AFLF_ can then be most conveniently realized with momentum-accelerated, gradient-following samplers such as Adam.^**46**^ The detailed AFLF configuration is listed in **Supplementary Table S3**. The AFLF dynamics are primarily driven by the repelling loss term, as shown in the loss profiles along the trajectory for each system (**Supplementary Figures S28, S29, S30, S31, S32, S33, S34**).

### 4.3 Computational Details

The AFLF program was implemented as an extension to the AF 2.3.2 source (https://github.com/google-deepmind/alphafold) based on the JAX 0.3.25 ecosystem (http://github.com/jax-ml/jax) and the Optax 0.1.7 library (https://github.com/google-deepmind/optax).

The AFLF checkpoints were retrieved from the default AF inference pipeline with 3 recycling iterations that embeds both the features and the positions from the previous prediction. The MSAs were resampled and clustered into 508 clusters and the structural templates were disabled. The AFLF inference modeling uses only the Evoformer and the StructMod blocks, and the original AF test-time configuration is thus obsolete. As the reverse-mode gradient computation in the Evoformer stack is memory-intensive, we executed the AFLF inference in the “jax.numpy.bfloat16” precision and rematerialized intermediate activations on demand during the backward sweep, such that the entire workload fit on a single 40 GB NVIDIA A40 GPU. The “model_1_ptm” set of model parameters was adopted for all systems. The per-conformation inference time ranges from a few seconds to approximately twenty seconds, depending on the sequence length and the checkpoint layer from which the latent inference is initiated (**Supplementary Table S4**).

For the reference MD simulations of ubiquitin with the CHARMM36m^**47**^ classical potential, the system was set up in CHARMM^**48**^ and production MD simulations were carried out with OpenMM^**49**^. The MDAnalysis,^**50**^ the MDpocket,^**51**^ the umap-learn,^**52**^ and the BioEmu-Benchmark^**13**^ libraries were applied for analysis. All molecular graphics were prepared with ChimeraX.^**53**^

## Acknowledgement

The authors thank Profs. Yi Qin Gao, Wei Yang, and Yaoqi Zhou for insightful discussions. This work is supported by the Shenzhen Medical Research Fund (B2404003), the National Natural Science Foundation of China (T2596084, 22503068), the Hangzhou Joint Fund of Zhejiang Provincial Natural Science Foundation of China (LHZQN26B030001), the Pioneer and Leading Goose R&D Program of Zhejiang (2023C03109), and the State Key Laboratory of Gene Expression (SKLGE-ZX-2026003). We thank the Westlake University Supercomputer Center for computational resources and related assistance.

## Author Contributions

**Runtong Qian**: Software, Validation, Formal analysis, Investigation, Data curation, Writing – review & editing, Visualization; **Rui Zhan**: Validation, Formal analysis, Investigation, Data curation, Writing – review & editing, Visualization; **Zilin Song**: Conceptualization, Methodology, Software, Validation, Formal analysis, Investigation, Data curation, Writing – original draft, Writing – review & editing, Visualization, Funding acquisition; **Jing Huang**: Conceptualization, Methodology, Validation, Formal analysis, Investigation, Resources, Writing – review & editing, Supervision, Project administration, Funding acquisition.

## Data Availability

Additional metadata are available in the Zenodo data repository with the identifier https://doi.org/10.5281/zenodo.23016771.

## Code Availability

The computer code developed in the current study is publicly accessible at the following repository: https://github.com/JingHuangLab/AFLF.git.

