## Supplemental Tables and Figures for "Zero-Shot Generation of Protein Conformational Ensembles Through AlphaFold Latent Flooding"

### 1 Supporting Methods

#### 1.1 Molecular Dynamics

Molecular dynamics simulations were used as the orthogonal reference for the structural fluctuations of the ubiquitin protein. The native folded structure (PDB id: 1UBQ) was solvated in a 70 Å cubic box. The CHARMM36m and the CHARMM-modified TIP3P force fields were used to model the protein and the solvent, respectively. The short-range nonbonded interactions were explicitly treated within 10 Å and were smoothened out at 12 Å. The periodic boundary conditions were employed in all three dimensions and the particle mesh Ewald summation was applied for long range electrostatic and dispersion interactions. All covalent bonds to hydrogen atoms were constrained. The solvated system was then subjected sequentially to an initial energy minimization, a 4 ns canonical (NVT) equilibration, and a 4 ns isothermal-isobaric (NPT) equilibration. The production run was carried out for 500 ns with temperature maintained at 300 K using the Nosé-Hoover chain thermostat and pressure coupled to 1 bar using the Monte Carlo barostat. The dynamical trajectory was integrated at 2 fs time steps and was recorded every 100 ps.

#### 1.2 Benchmark Evaluations

We selected nine proteins with experimental $B$-factors and generated the AFLF ensembles using the ubiquitin protocol described in the main text, with all settings held identical across the nine proteins. For each protein, the AFLF ensemble was generated as four independent 5,000-frame trajectories initiated from the cached Evoformer checkpoint of the native AF prediction. The reference ensembles of BioEmu and AFsample2 were generated in this work using the released models, with four independent batches of 5,000 conformations collected for each method. The $B$-factors were computed from the root mean square fluctuations as described in the main text.

We further evaluated AFLF on the three benchmark sets compiled in the BioEmu study, namely the 44 domain-motion systems, the 34 cryptic-pocket systems, and the 38 OOD-60 systems. For each system, the AFLF ensemble was generated as five independent 5,000-frame trajectories initiated from the cached checkpoint at the 40th Evoformer layer of the native AF prediction. The BioEmu reference ensembles were taken from the publicly released benchmark data and analyzed together with the AFLF ensembles using the native benchmark metrics implemented in BioEmu-Benchmark. All other AFLF parameters were held identical within each set and were inherited from the corresponding prototype system described in the main text: the domain-motion and the OOD-60 sets adopted the adenylate kinase protocol, and the cryptic-pocket set adopted the TEM-1 protocol (**Supplementary Table S3**). $0.1\%$ coverage was used as metric, which is the fraction of reference conformations matched below an RMSD threshold by at least one of 1,000 samples drawn with replacement, averaged over 20 bootstrap replicates. Global and local backbone RMSD were used for domain motions and cryptic pocket formation, respectively.

#### 1.3 Latent Visualizations

The latent dynamics driven by AFLF can be quantified by the evolution of the massive activations (the top and bottom $1\%$ tensor elements) in the latent states. We computed the set overlap of the massive activations by the Jaccard Index (JI). Formally, let $\mathbf{B_h}(\tau)$ be the binary matrix of identical dimensions that flags the massive activations and $\mathcal{B}_\mathbf{h}(\tau)$ the subset of linearized indices of the nonzero entries in $\mathbf{B_h}(\tau)$, the JI is defined,

$$\text{JI}(\tau, \tau') := \frac{\#\left(\mathcal{B}_\mathbf{h}(\tau) \cap \mathcal{B}_\mathbf{h}(\tau')\right)}{\#\left(\mathcal{B}_\mathbf{h}(\tau) \cup \mathcal{B}_\mathbf{h}(\tau')\right)}, \tag{1}$$

such that it quantifies the overlap of massively activated entries between latent states at different times. The UMAP dimensionality reduction was performed by using the pairwise JIs as the distance metric. The optimal transport distance in one dimension quantifies the cost of mutating two balanced matrices into each other and is obtained by the monotone coupling,

$$d_{\text{OT}}(\tau, \tau') := \frac{1}{\sum \mathbf{B_h}(\tau)} \sum \left\| \text{sort}(\mathbf{B_h}(\tau)\mathbf{h}(\tau)) - \text{sort}(\mathbf{B_h}(\tau')\mathbf{h}(\tau')) \right\|_1 . \tag{2}$$

Empirically, we used JI to monitor the dynamical evolution in positions of the massive activations and $d_{\mathrm{OT}}$ in value distributions.

#### 2 Supporting Figures

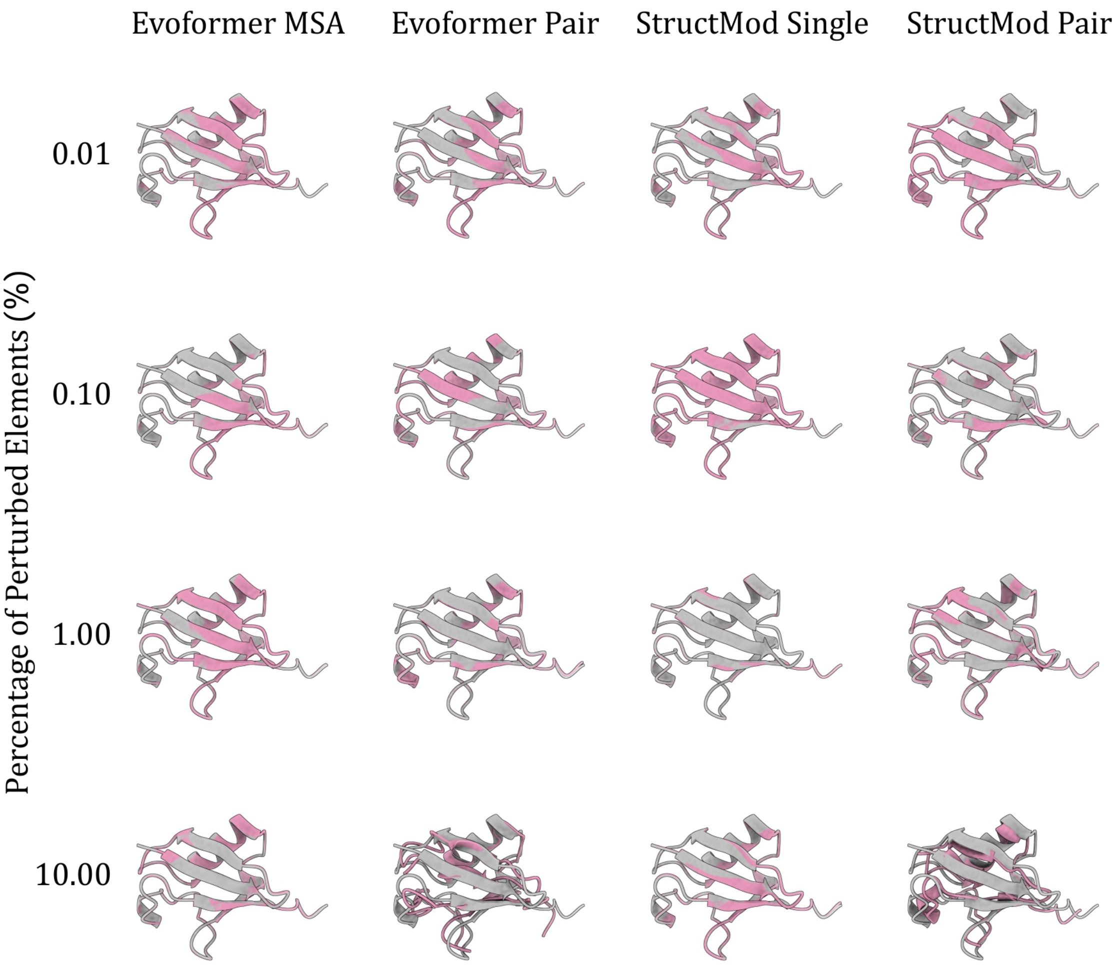

**Supplementary Figure S1.** Ubiquitin structures predicted from scheme0 latent state perturbations. The structure predicted from perturbed latent states (pink) are superposed with the unperturbed predictions (gray). All structures were rendered at the same scale.

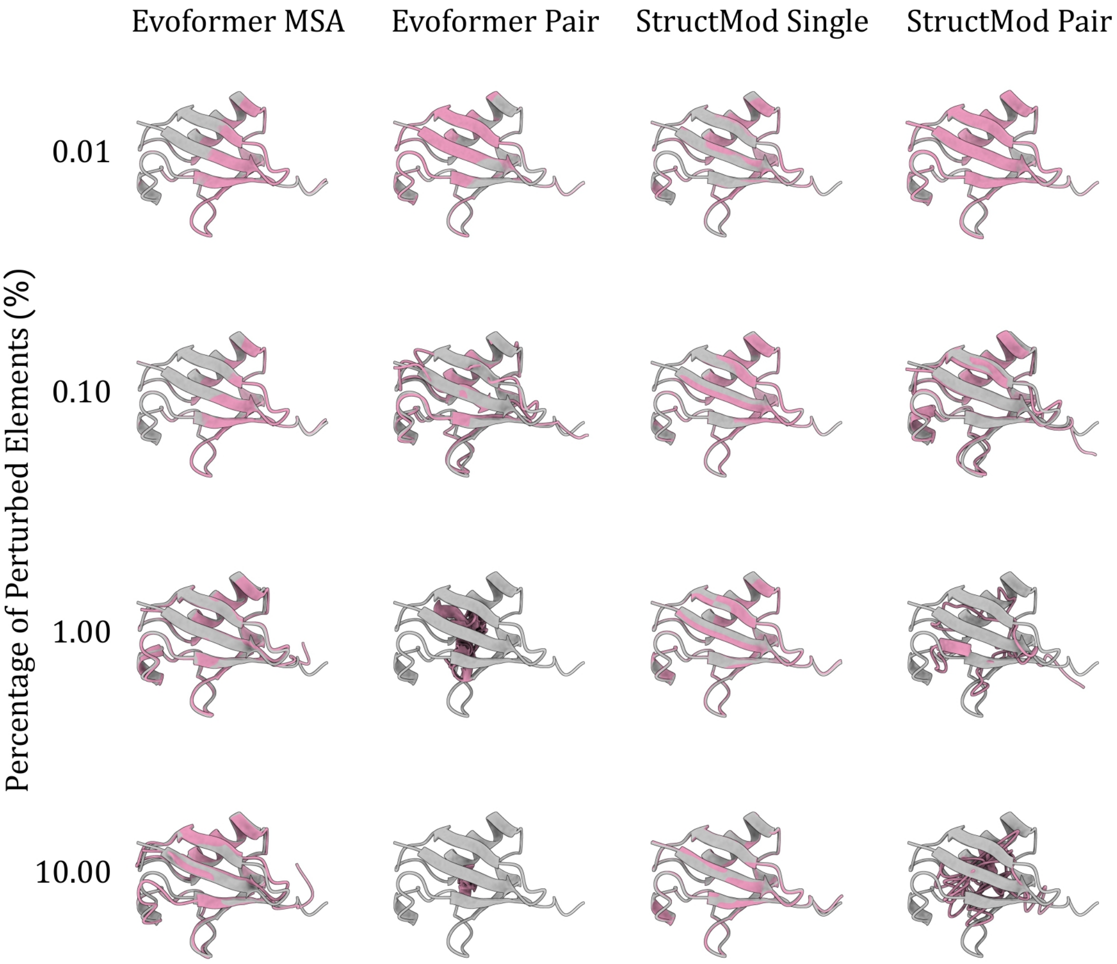

**Supplementary Figure S2.** Ubiquitin structures predicted from scheme1 latent state perturbations. The structure predicted from perturbed latent states (pink) are superposed with the unperturbed predictions (gray). All structures were rendered at the same scale.

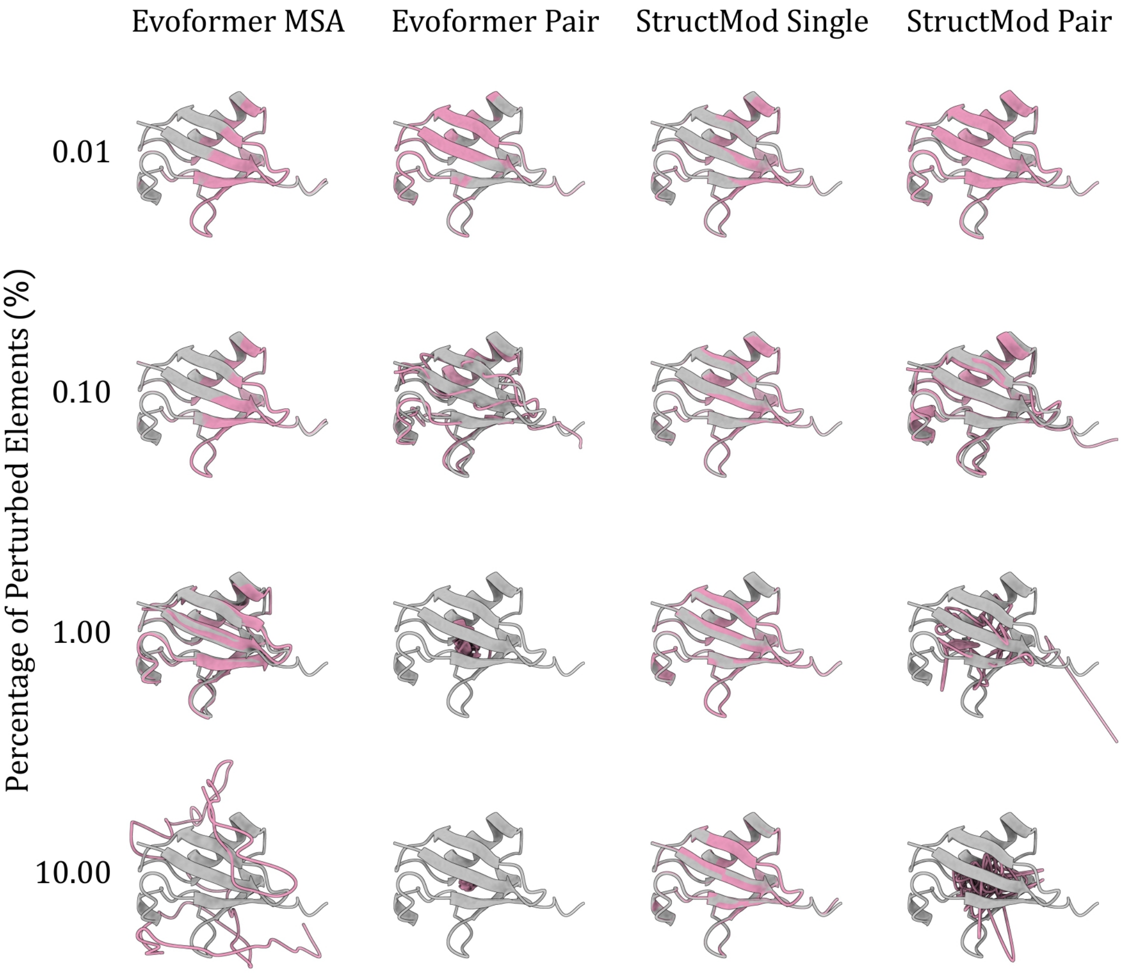

**Supplementary Figure S3.** Ubiquitin structures predicted from scheme2 latent state perturbations. The structure predicted from perturbed latent states (pink) are superposed with the unperturbed predictions (gray). All structures were rendered at the same scale.

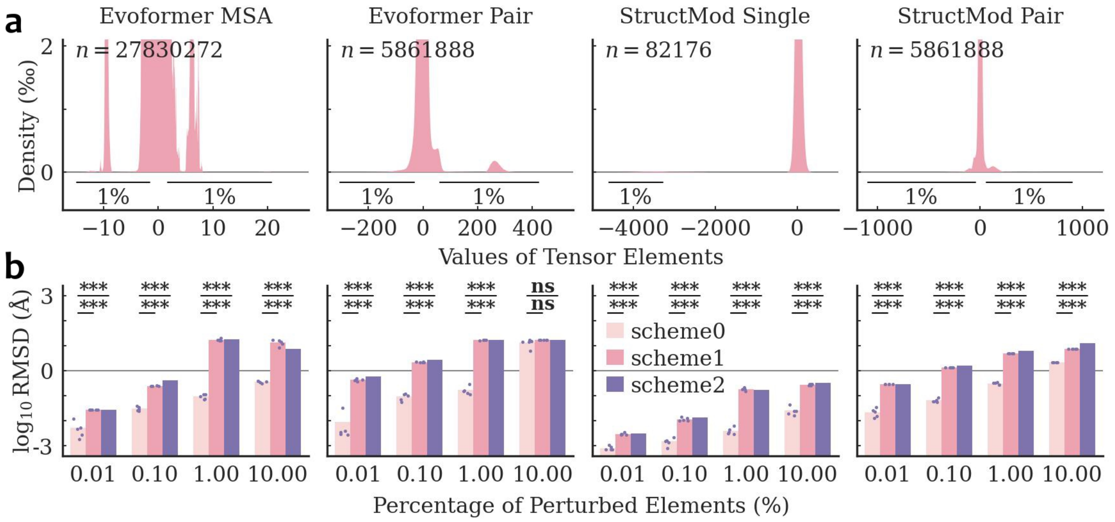

**Supplementary Figure S4.** Latent features distribution and adenylate kinase (AdK) structural deviations from perturbed latent features. **(a)** Permillage value distributions of the AF2 latent tensor elements used to predict AdK conformations. The 1st and 99th percentiles are indicated and $n$ is the total number of elements in each tensor. **(b)** Backbone root mean squared deviations ($\log_{10}$RMSDs) of AdK conformations predicted from perturbed latent tensors relative to the native unperturbed prediction. Statistical tests: scheme0 ($n = 5$) versus scheme1 ($n = 5$) by independent samples $t$-test; scheme0 ($n = 5$) versus scheme2 ($n = 1$) by one-sample $t$-test. ***: $p < 0.001$; **: $p < 0.01$; *: $p < 0.05$; **ns**: not significant.

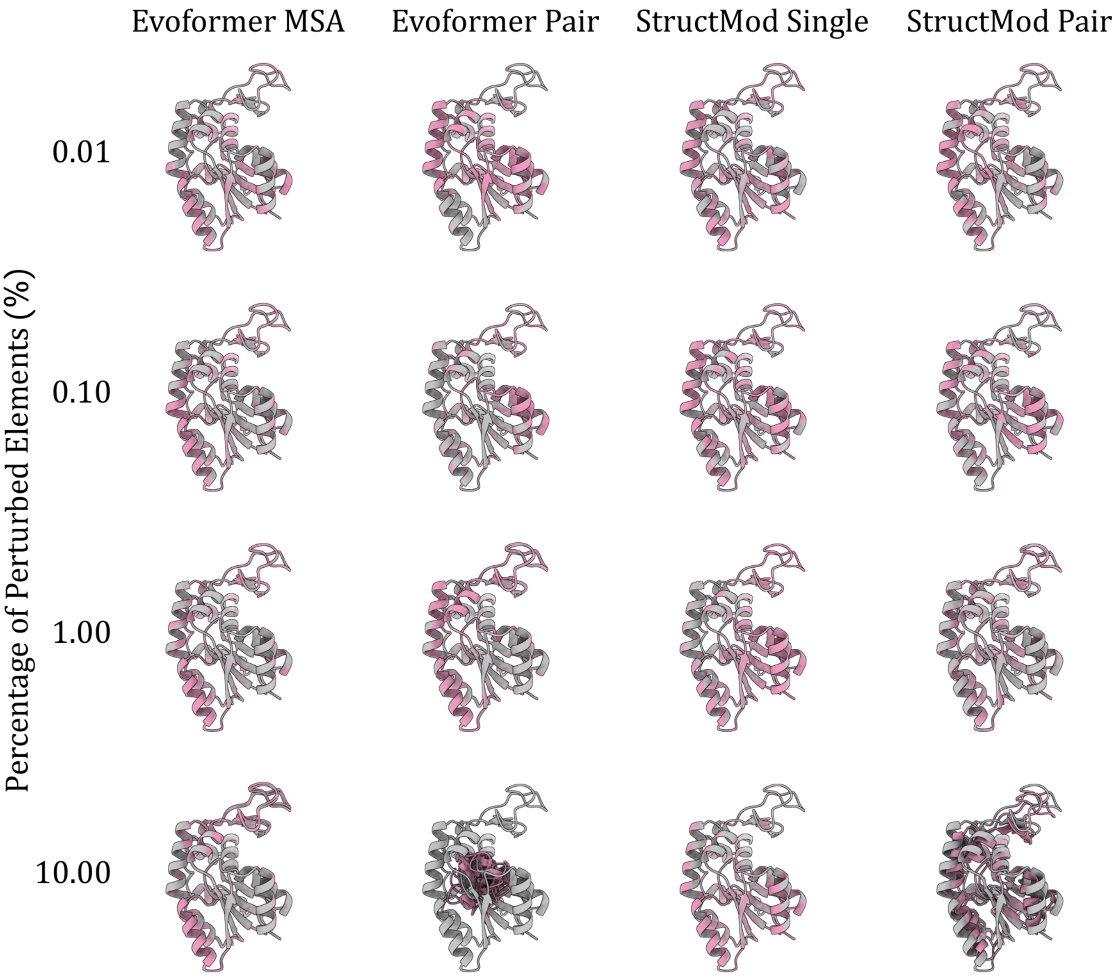

**Supplementary Figure S5.** Adenylate kinase structures predicted from scheme0 latent state perturbations. The structure predicted from perturbed latent states (pink) are superposed with the unperturbed predictions (gray). All structures were rendered at the same scale.

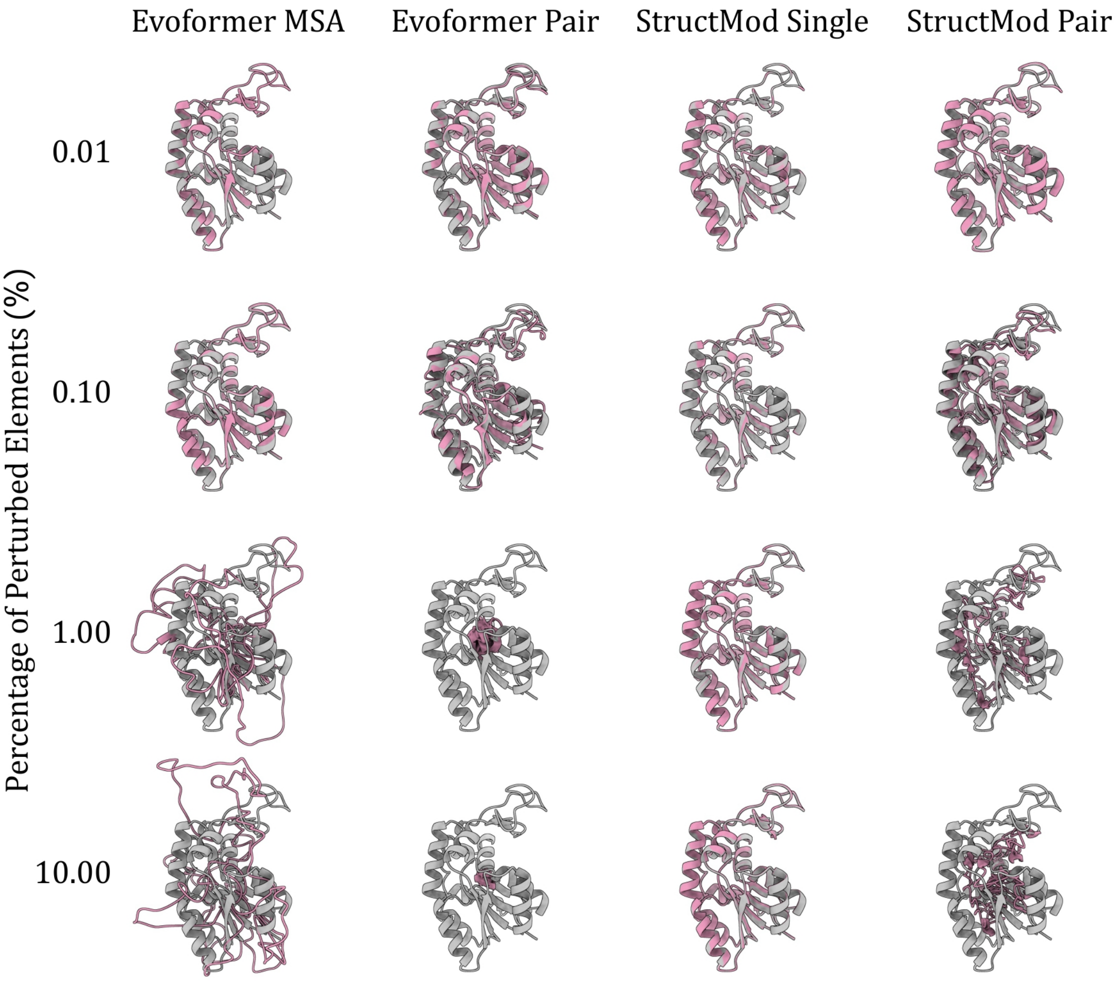

**Supplementary Figure S6.** Adenylate kinase structures predicted from scheme1 latent state perturbations. The structure predicted from perturbed latent states (pink) are superposed with the unperturbed predictions (gray). All structures were rendered at the same scale.

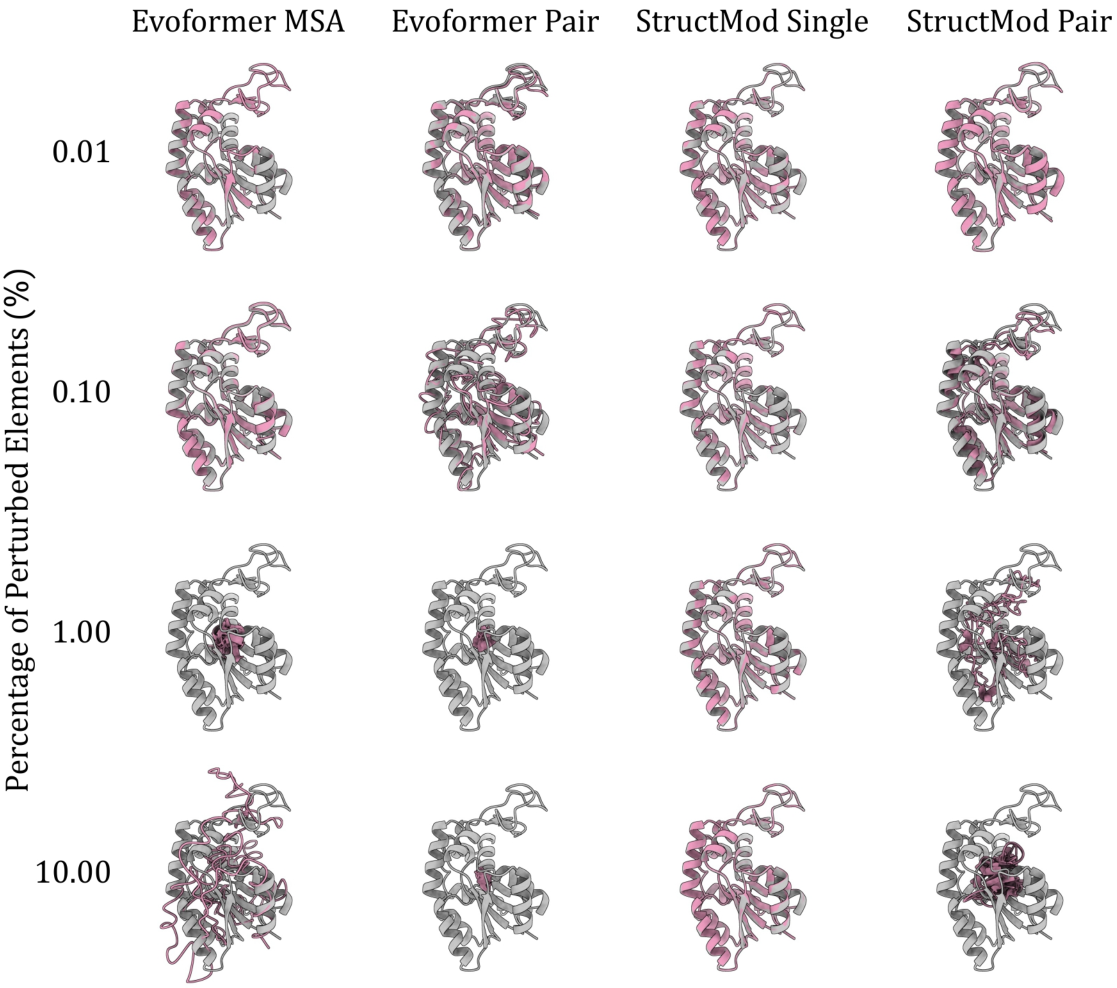

**Supplementary Figure S7.** Adenylate kinase structures predicted from scheme2 latent state perturbations. The structure predicted from perturbed latent states (pink) are superposed with the unperturbed predictions (gray). All structures were rendered at the same scale.

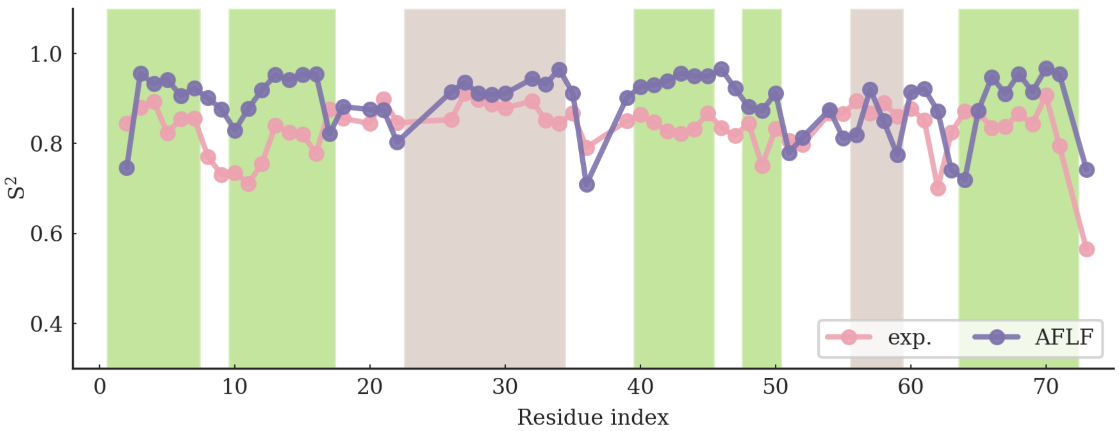

**Supplementary Figure S8.** The backbone N–H order parameters ($S^2$) of ubiquitin from NMR relaxation and from the AFLF ensemble, with the N–H vectors of the generated conformations assigned from the backbone heavy atoms. The secondary structures are annotated (green for extended strands and brown for helices).

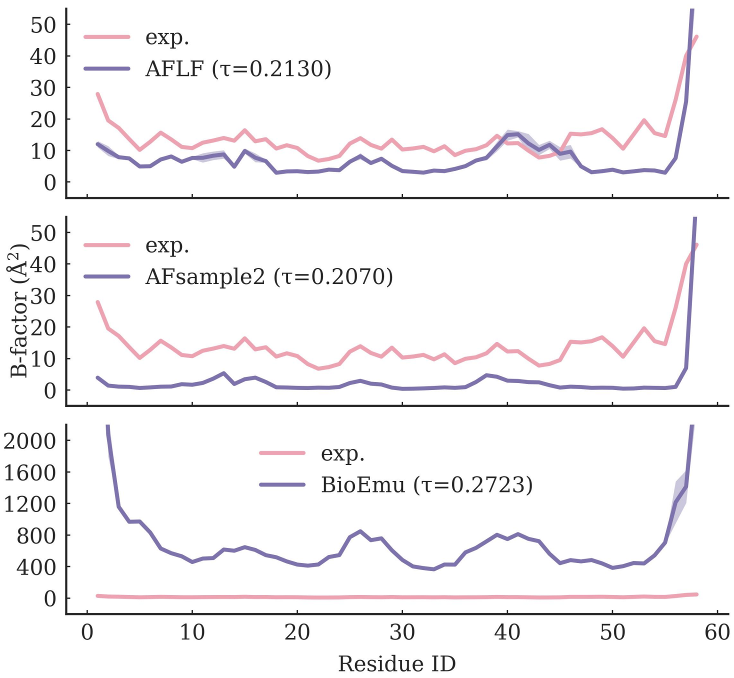

**Supplementary Figure S9.** Per-residue $B$-factor profiles of bovine pancreatic trypsin inhibitor, BPTI (PDB id: 5PTI) from experiment (pink) and simulations (purple): AFLF (top), AFsample2 (middle), and BioEmu (bottom). For each method, the $B$-factors were derived from the RMSF computed over 20,000 frames formed by concatenating four independent 5,000-frame trajectories and the shaded bands indicate the standard deviation across the four trajectories. The Kendall $\tau$ coefficient against the experimental profile is annotated in each panel. Note the different ordinate scale of the BioEmu panel.

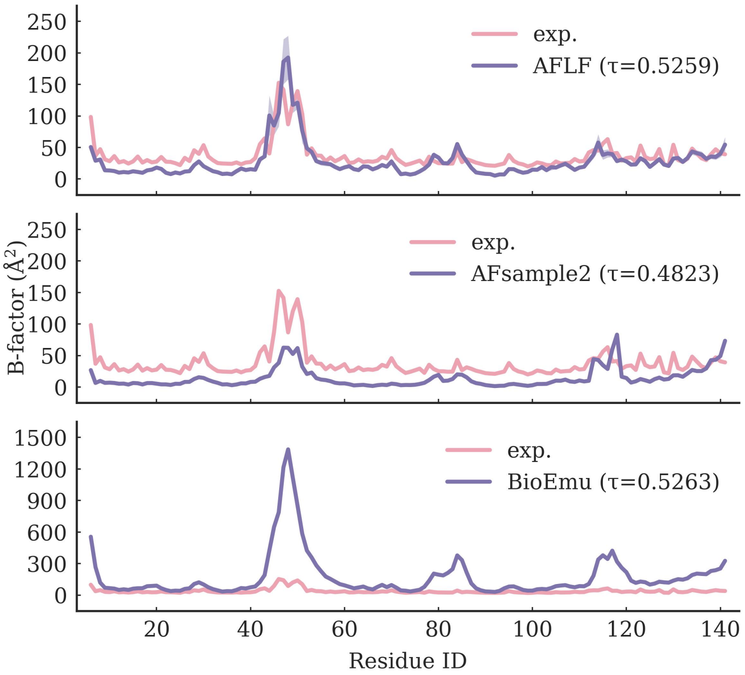

**Supplementary Figure S10.** Per-residue $B$-factor profiles of P00644 (PDB id: 1EY0) from experiment (pink) and simulations (purple): AFLF (top), AFsample2 (middle), and BioEmu (bottom). For each method, the $B$-factors were derived from the RMSF computed over 20,000 frames formed by concatenating four independent 5,000-frame trajectories and the shaded bands indicate the standard deviation across the four trajectories. The Kendall $\tau$ coefficient against the experimental profile is annotated in each panel. Note the different ordinate scale of the BioEmu panel.

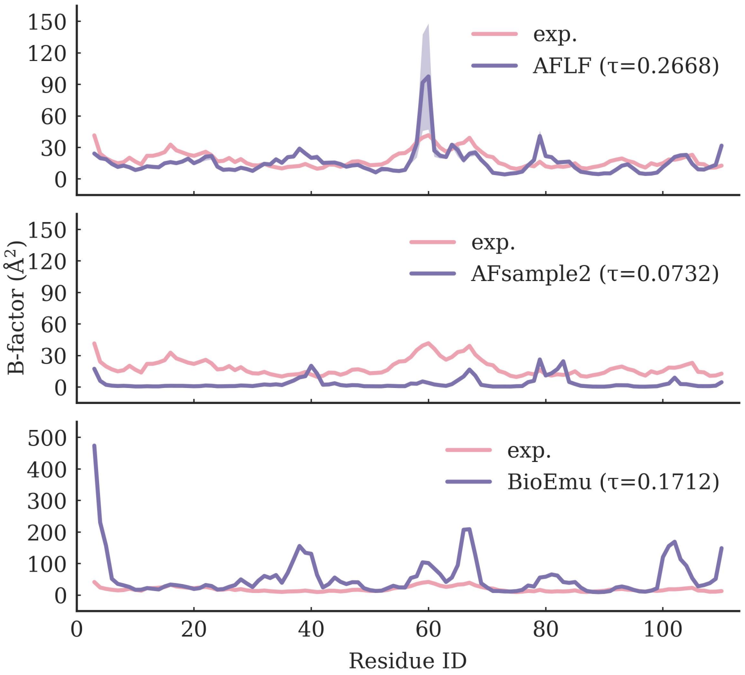

**Supplementary Figure S11.** Per-residue $B$-factor profiles of P00648 (PDB id: 1A2P) from experiment (pink) and simulations (purple): AFLF (top), AFsample2 (middle), and BioEmu (bottom). For each method, the $B$-factors were derived from the RMSF computed over 20,000 frames formed by concatenating four independent 5,000-frame trajectories and the shaded bands indicate the standard deviation across the four trajectories. The Kendall $\tau$ coefficient against the experimental profile is annotated in each panel. Note the different ordinate scale of the BioEmu panel.

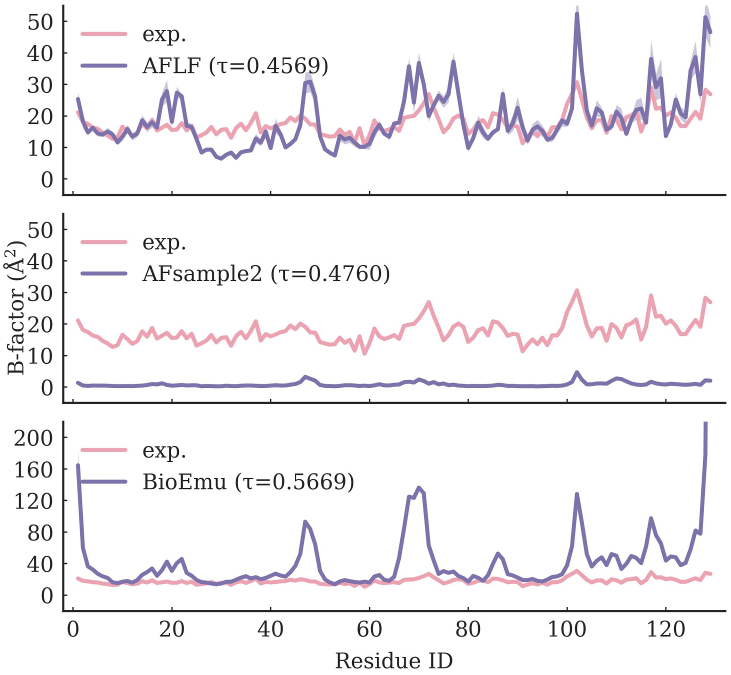

**Supplementary Figure S12.** Per-residue $B$-factor profiles of P00698 (PDB id: 1AKI) from experiment (pink) and simulations (purple): AFLF (top), AFsample2 (middle), and BioEmu (bottom). For each method, the $B$-factors were derived from the RMSF computed over 20,000 frames formed by concatenating four independent 5,000-frame trajectories and the shaded bands indicate the standard deviation across the four trajectories. The Kendall $\tau$ coefficient against the experimental profile is annotated in each panel. Note the different ordinate scale of the BioEmu panel.

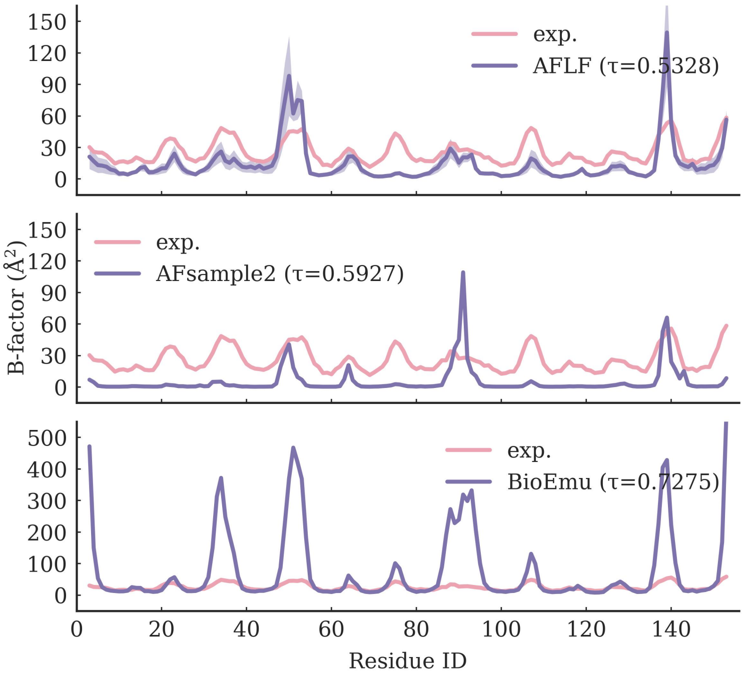

**Supplementary Figure S13.** Per-residue $B$-factor profiles of P01854 (PDB id: 1I1B) from experiment (pink) and simulations (purple): AFLF (top), AFsample2 (middle), and BioEmu (bottom). For each method, the $B$-factors were derived from the RMSF computed over 20,000 frames formed by concatenating four independent 5,000-frame trajectories and the shaded bands indicate the standard deviation across the four trajectories. The Kendall $\tau$ coefficient against the experimental profile is annotated in each panel. Note the different ordinate scale of the BioEmu panel.

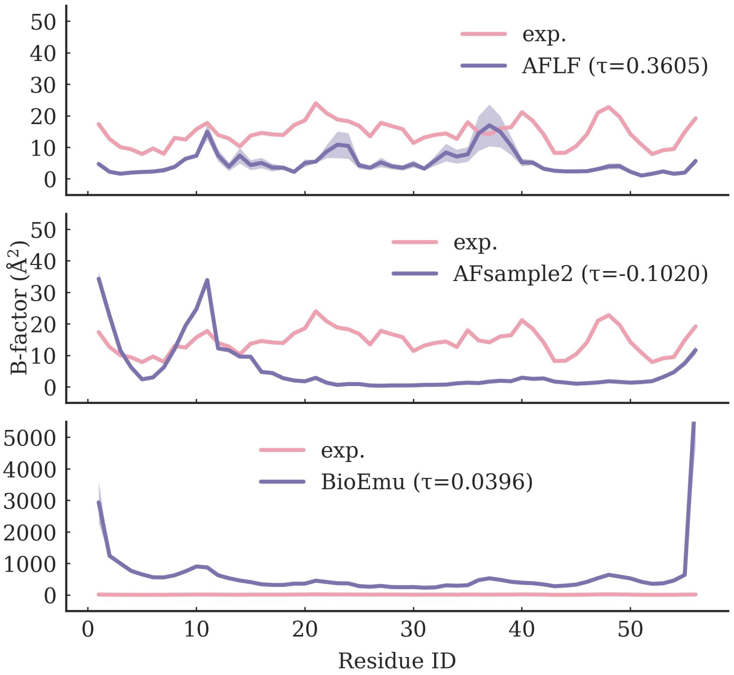

**Supplementary Figure S14.** Per-residue $B$-factor profiles of P06654 (PDB id: 1PGB) from experiment (pink) and simulations (purple): AFLF (top), AFsample2 (middle), and BioEmu (bottom). For each method, the $B$-factors were derived from the RMSF computed over 20,000 frames formed by concatenating four independent 5,000-frame trajectories and the shaded bands indicate the standard deviation across the four trajectories. The Kendall $\tau$ coefficient against the experimental profile is annotated in each panel. Note the different ordinate scale of the BioEmu panel.

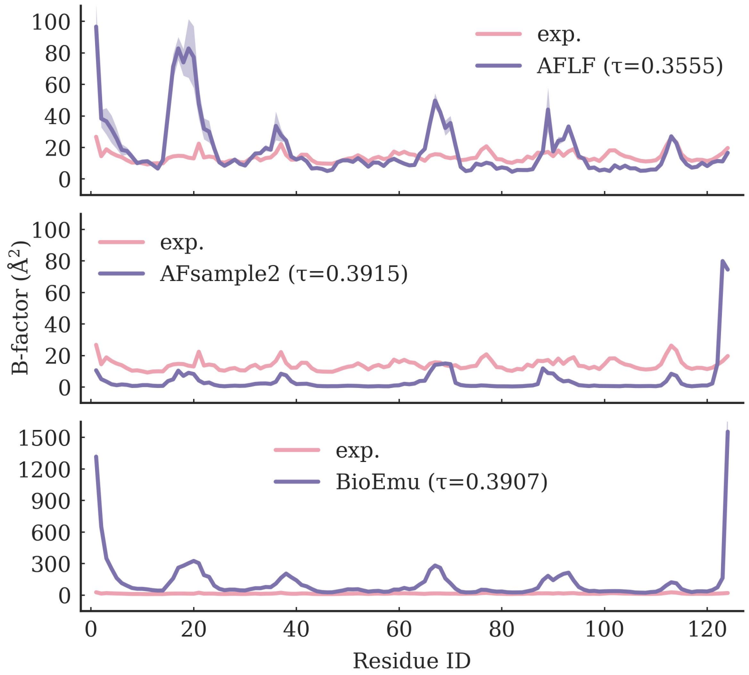

**Supplementary Figure S15.** Per-residue $B$-factor profiles of P61823 (PDB id: 1KF5) from experiment (pink) and simulations (purple): AFLF (top), AFsample2 (middle), and BioEmu (bottom). For each method, the $B$-factors were derived from the RMSF computed over 20,000 frames formed by concatenating four independent 5,000-frame trajectories and the shaded bands indicate the standard deviation across the four trajectories. The Kendall $\tau$ coefficient against the experimental profile is annotated in each panel. Note the different ordinate scale of the BioEmu panel.

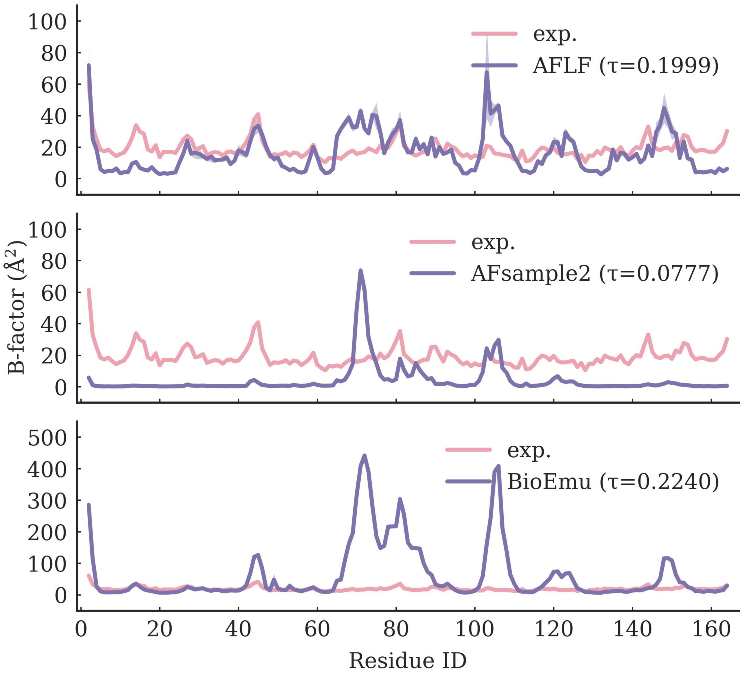

**Supplementary Figure S16.** Per-residue $B$-factor profiles of P62937 (PDB id: 3K0N) from experiment (pink) and simulations (purple): AFLF (top), AFsample2 (middle), and BioEmu (bottom). For each method, the $B$-factors were derived from the RMSF computed over 20,000 frames formed by concatenating four independent 5,000-frame trajectories and the shaded bands indicate the standard deviation across the four trajectories. The Kendall $\tau$ coefficient against the experimental profile is annotated in each panel. Note the different ordinate scale of the BioEmu panel.

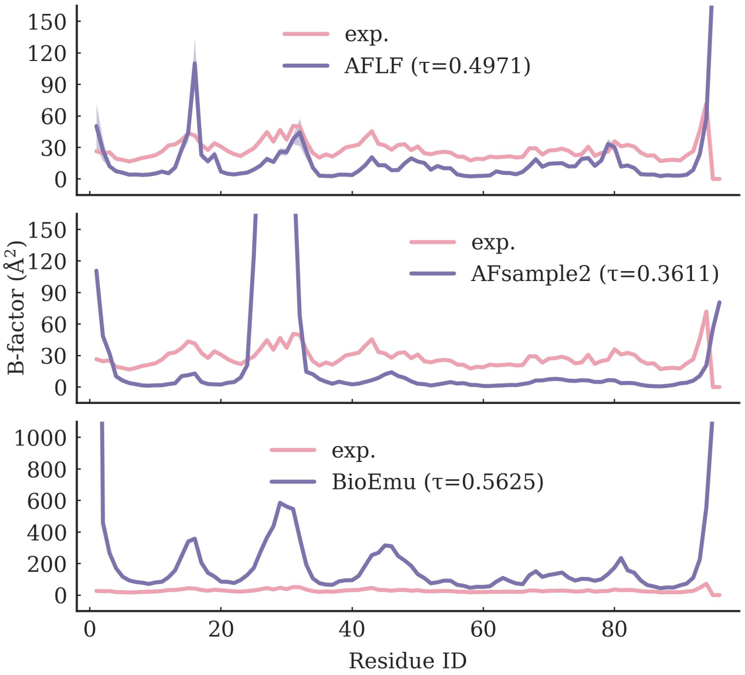

**Supplementary Figure S17.** Per-residue $B$-factor profiles of second PSD95/Discs large/ZO-1 domain, PDZ2 (PDB id: 3LNX) from experiment (pink) and simulations (purple): AFLF (top), AFsample2 (middle), and BioEmu (bottom). For each method, the $B$-factors were derived from the RMSF computed over 20,000 frames formed by concatenating four independent 5,000-frame trajectories and the shaded bands indicate the standard deviation across the four trajectories. The Kendall $\tau$ coefficient against the experimental profile is annotated in each panel. Note the different ordinate scale of the BioEmu panel.

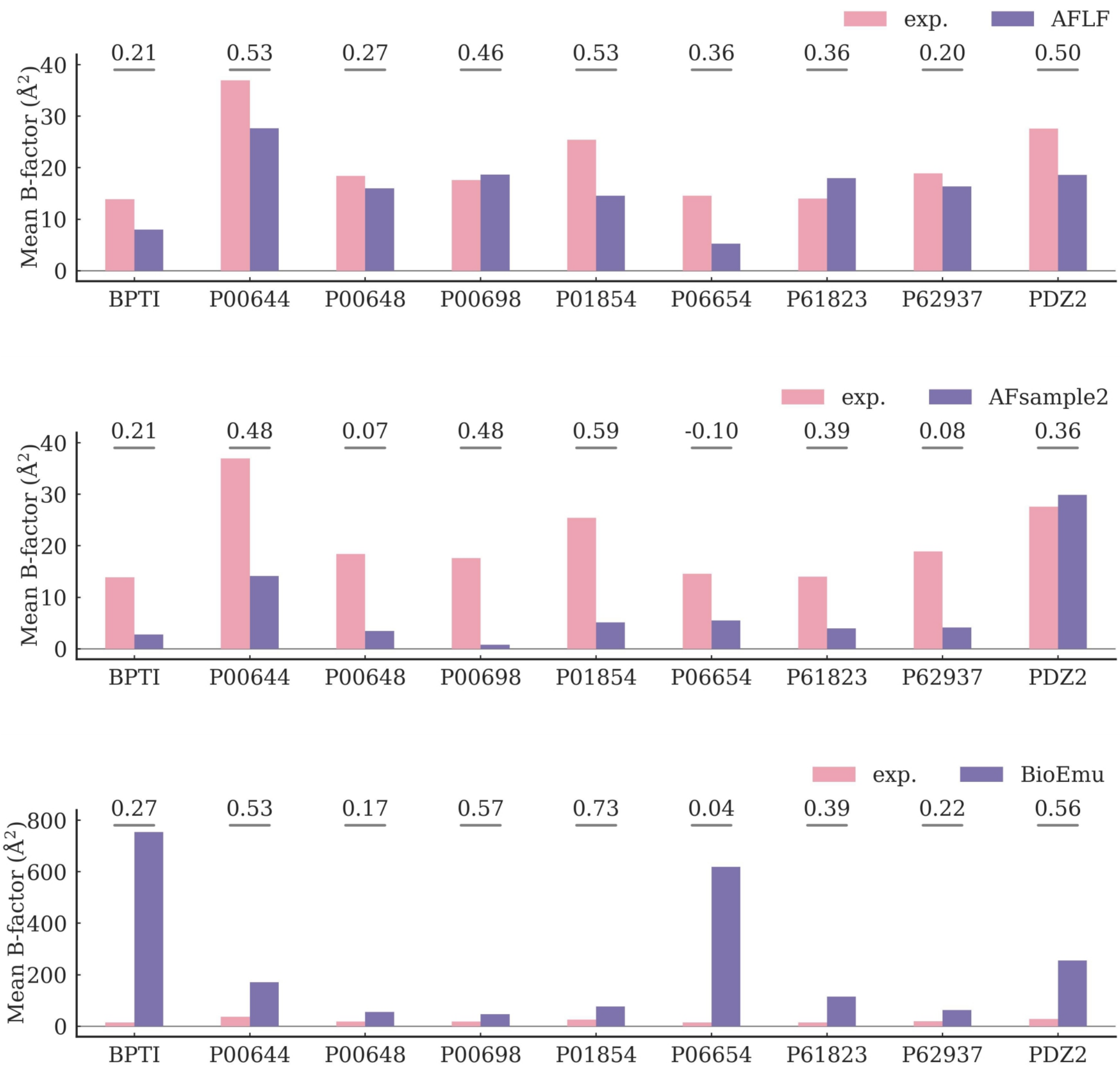

**Supplementary Figure S18.** Mean $B$-factors of the nine test proteins from experiment (pink) and simulations (purple): AFLF (top), AFsample2 (middle), and BioEmu (bottom). For each method, the $B$-factors were derived from the RMSF computed over 20,000 frames formed by concatenating four independent 5,000-frame trajectories, and the Kendall $\tau$ coefficients against the experimental per-residue profiles are annotated above the bars. Note the different ordinate scale of the BioEmu panel.

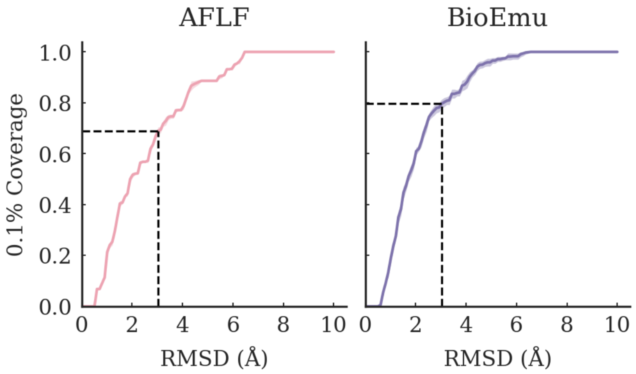

**Supplementary Figure S19.** The 0.1% coverage of the experimental alternative states on the 44 systems from the BioEmu domain-motion benchmark. The coverages at 3.0 Å are 0.690 for AFLF and 0.797 for BioEmu (dashed lines).

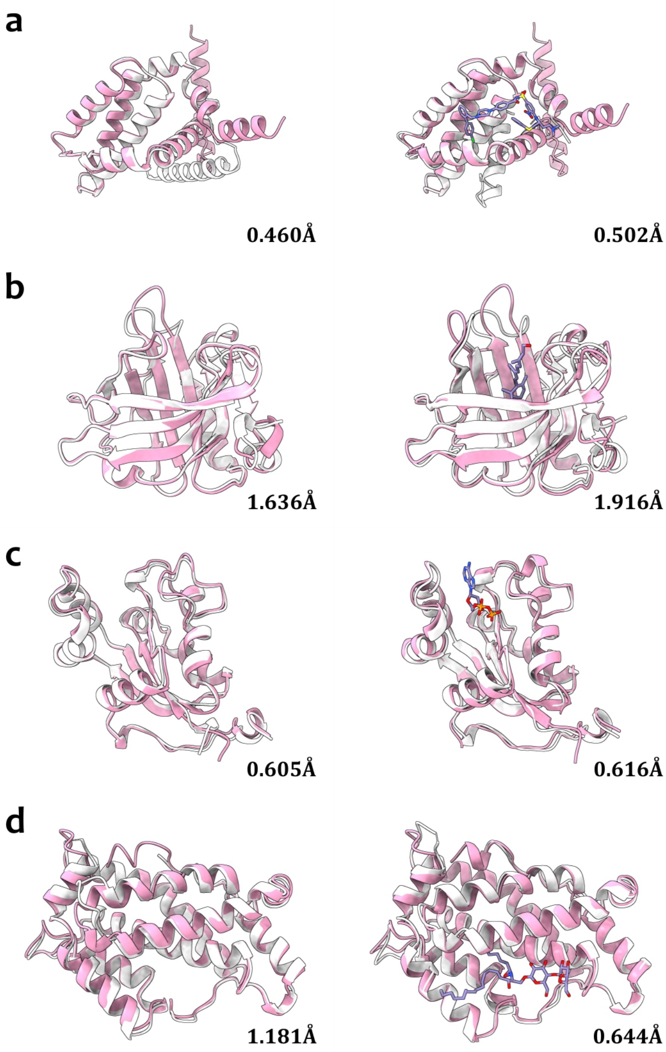

**Supplementary Figure S20.** The representative AFLF generated structures for the cryptic binding sites. **(a)** The anti-apoptotic protein Bcl-x$_L$; **(b)** The human nucleoside diphosphate kinase A (NDPK-A); **(c)** The glycolipid transfer protein (GLTP). The minimum C$\alpha$ RMSDs between the AFLF generated conformations (pink) and the experimental crystal structures (proteins in gray and ligands in purple) are listed.

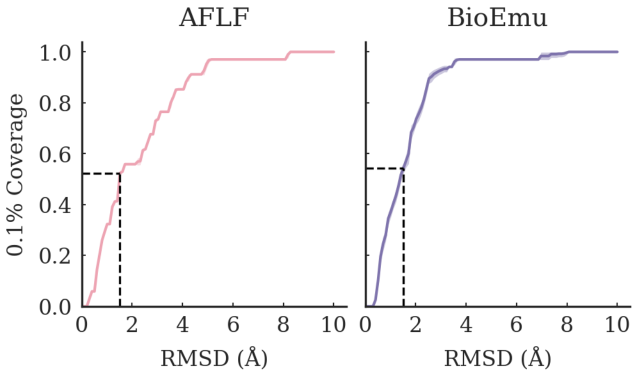

**Supplementary Figure S21.** The 0.1% coverage of the apo states on the 34 systems from the BioEmu cryptic-pocket benchmark. The coverages at 1.5 Å are 0.524 for AFLF and 0.543 for BioEmu (dashed lines).

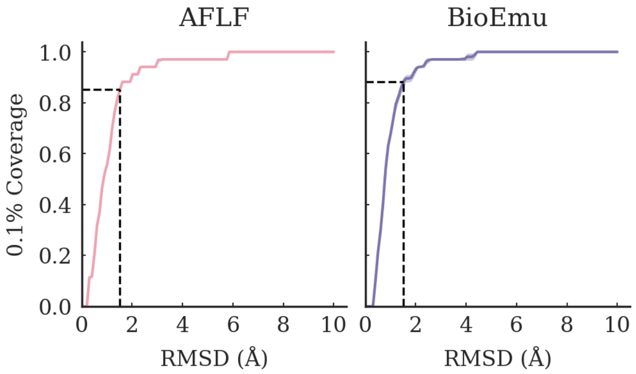

**Supplementary Figure S22.** The 0.1% coverage of the holo states on the 34 systems from the BioEmu cryptic-pocket benchmark. The coverages at 1.5 Å are 0.851 for AFLF and 0.882 for BioEmu (dashed lines).

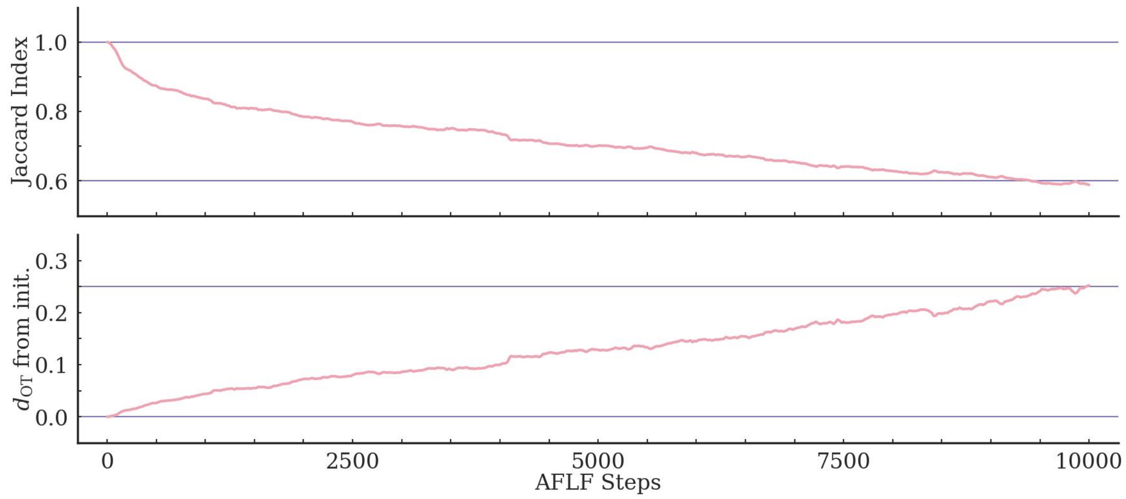

**Supplementary Figure S23.** The evolution of the Jaccard indices (top) and the optimal transport distances (bottom) with regard to the initial state along the adenylate kinase (AdK) trajectory.

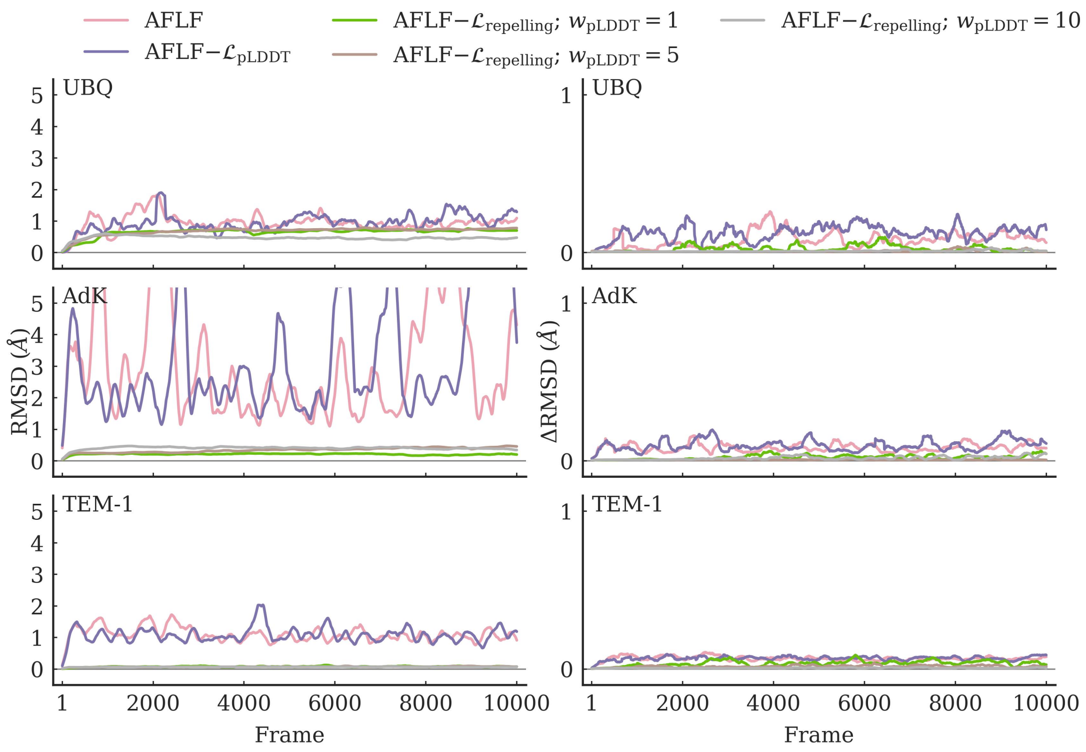

**Supplementary Figure S24.** Conformational exploration along the trajectories of the full and the ablated AFLF objectives. The backbone RMSD from the initial structure (left) and the per-frame $\Delta$RMSD (right) are shown for ubiquitin, AdK, and TEM-1 under the full AFLF objective, the objective without the pLDDT term, and the objective without the repelling term at the confidence weights $w_{\text{pLDDT}} = 1$, $5$, and $10$. The traces were smoothed by a sliding-window average over 200 consecutive frames.

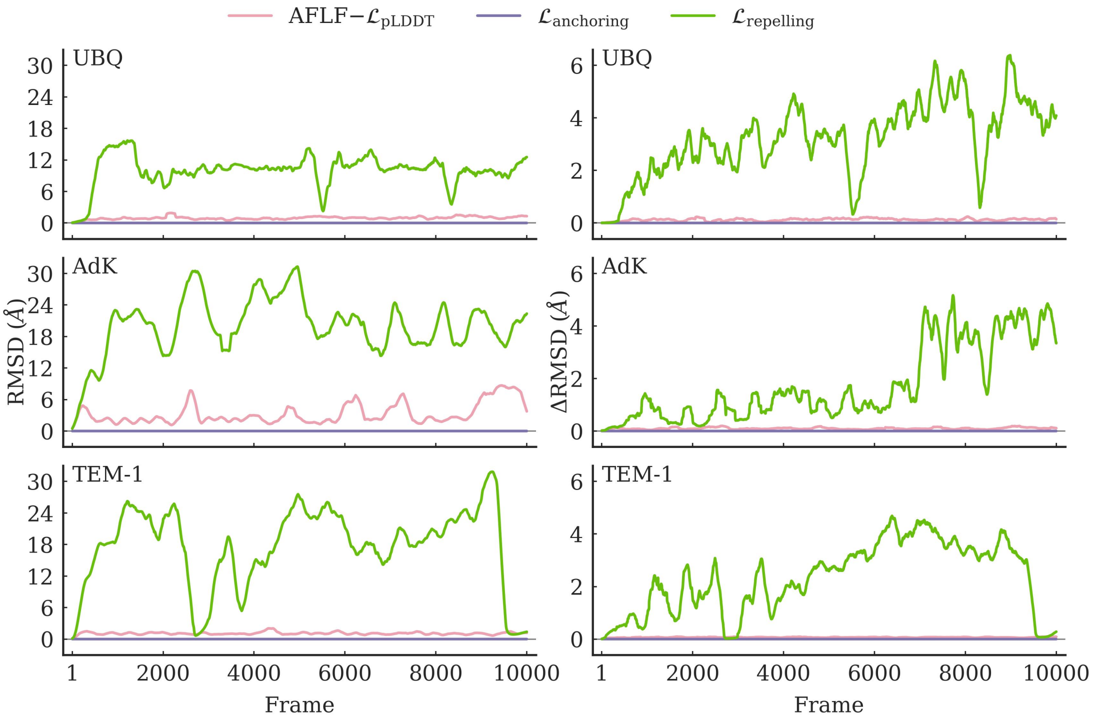

**Supplementary Figure S25.** Conformational exploration under the single-term objectives. The backbone RMSD from the initial structure (left) and the per-frame ΔRMSD (right) are shown for ubiquitin, AdK, and TEM-1 under AFLF−$\mathcal{L}_{\text{pLDDT}}$ and under the objectives retaining only the anchoring term ($\mathcal{L}_{\text{anchoring}}$) or only the repelling term ($\mathcal{L}_{\text{repelling}}$), with all other loss terms disabled. The anchoring-only objective keeps the trajectory at the initial structure, whereas the repelling-only objective drives large and uncontrolled structural deviations. The traces were smoothed by a sliding-window average over 200 consecutive frames.

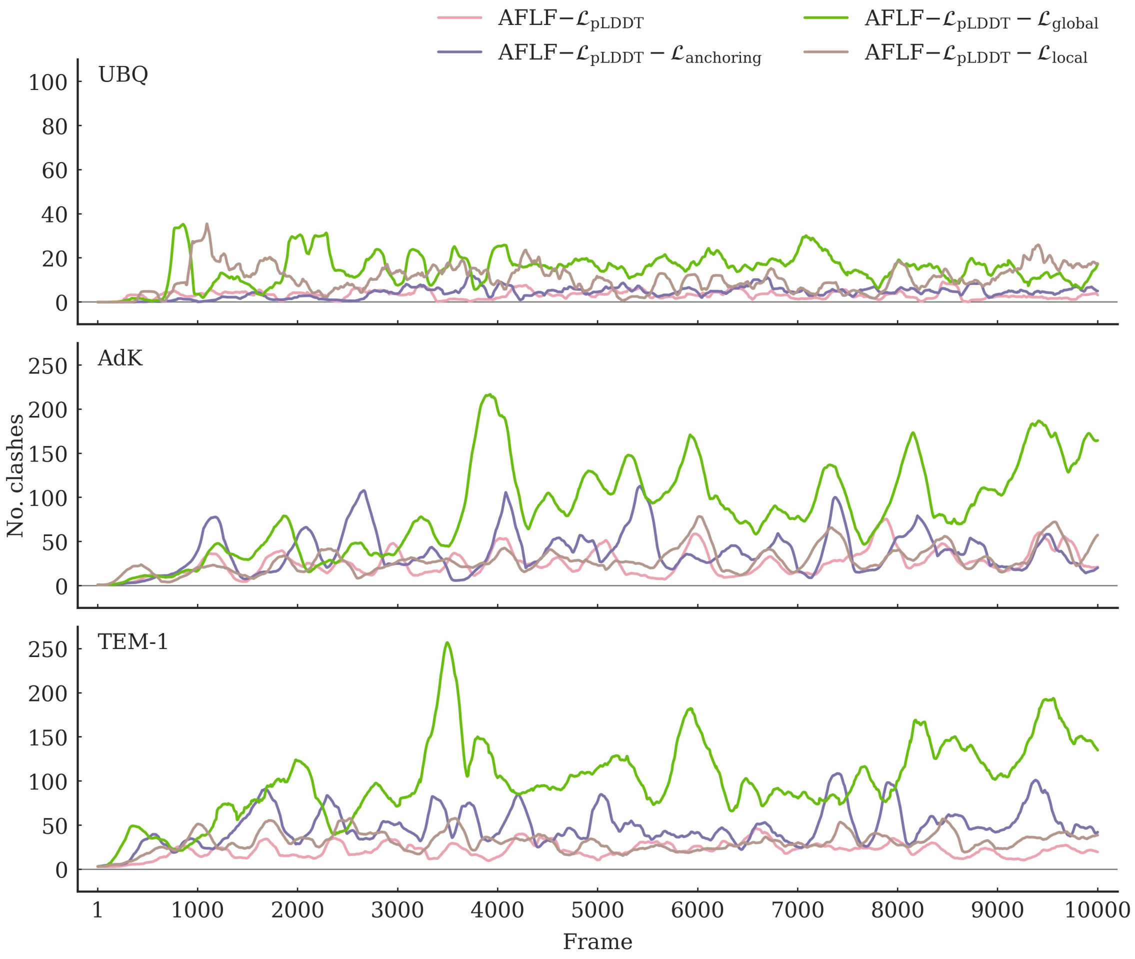

**Supplementary Figure S26.** Steric clashes along the trajectories of the ablated AFLF variants. The per-frame numbers of steric clashes are shown for ubiquitin, AdK, and TEM-1 under AFLF−$\mathcal{L}_{\text{pLDDT}}$ and under the variants with the anchoring, the global, or the local term additionally disabled. A steric clash was counted for each pair of heavy atoms closer than 2 Å, excluding pairs within the same residue or between residues adjacent in sequence. The traces were smoothed by a sliding-window average over 200 consecutive frames.

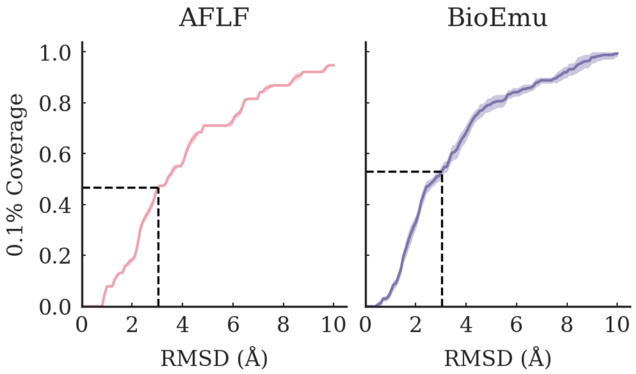

**Supplementary Figure S27.** The 0.1% coverage of the experimental alternative states on the 38 systems from the BioEmu OOD-60 benchmark. The coverages at 3.0 Å are 0.468 for AFLF and 0.530 for BioEmu (dashed lines).

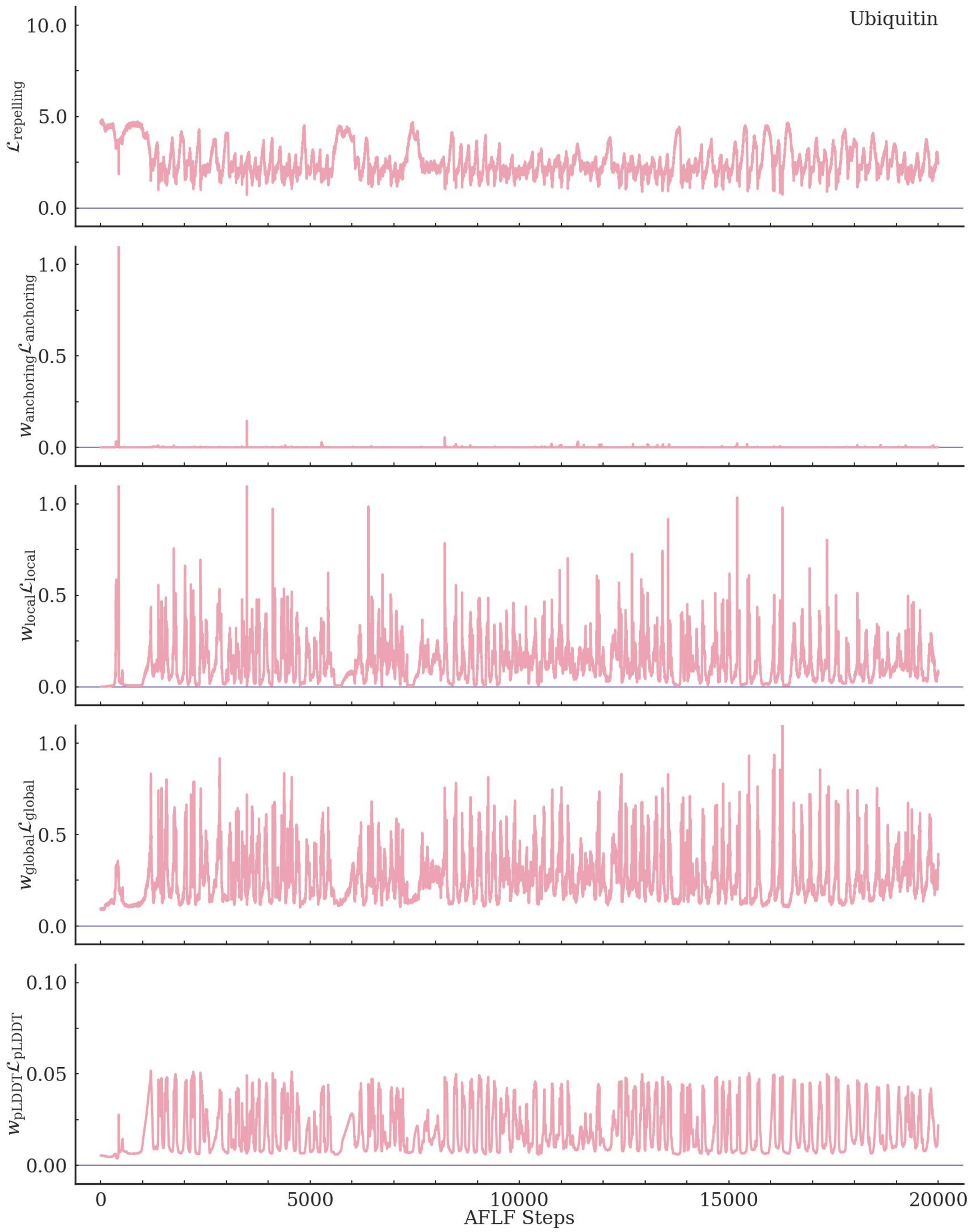

**Supplementary Figure S28.** The evolution of the loss terms along the ubiquitin trajectory.

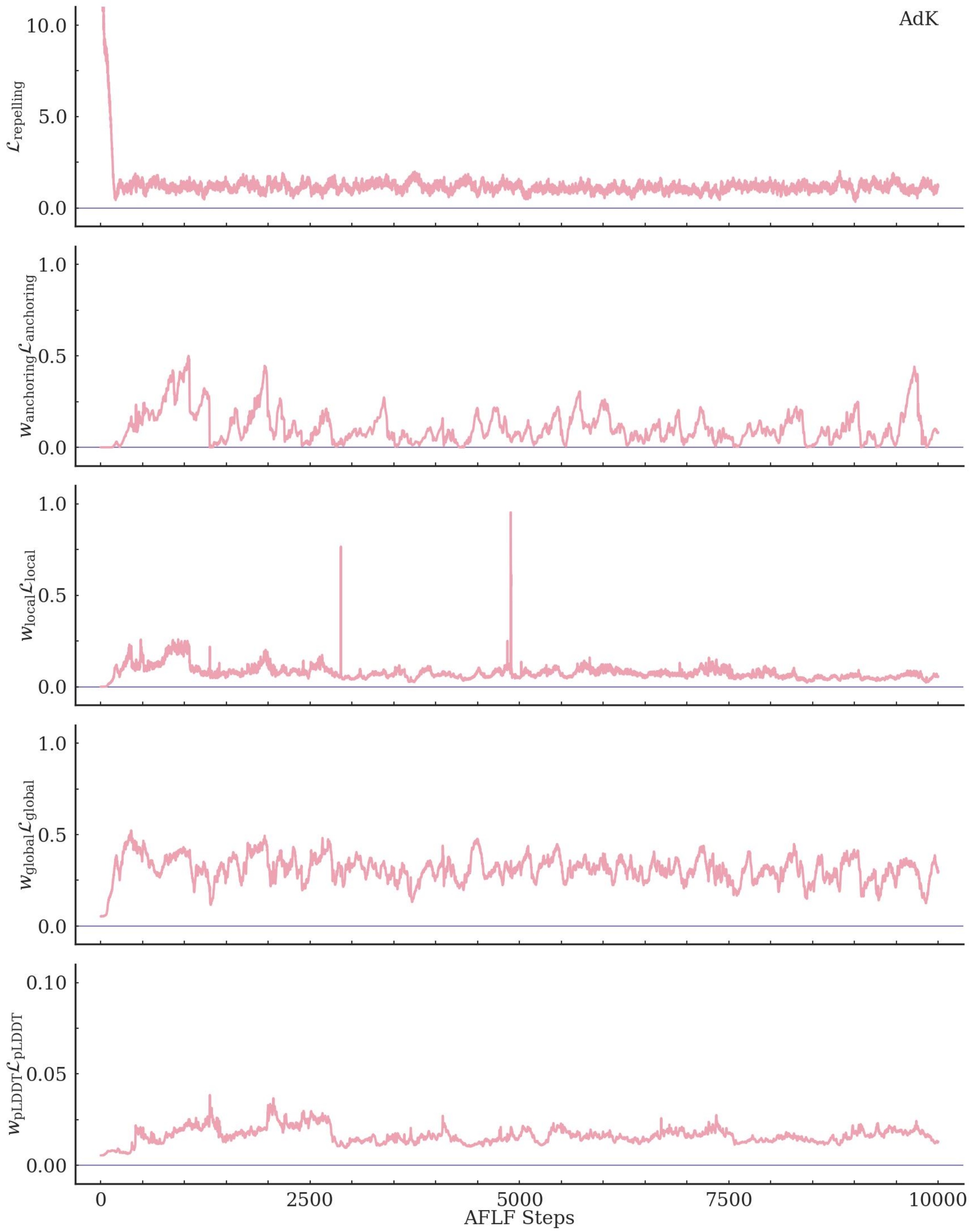

**Supplementary Figure S29.** The evolution of the loss terms along the adenylate kinase (AdK) trajectory.

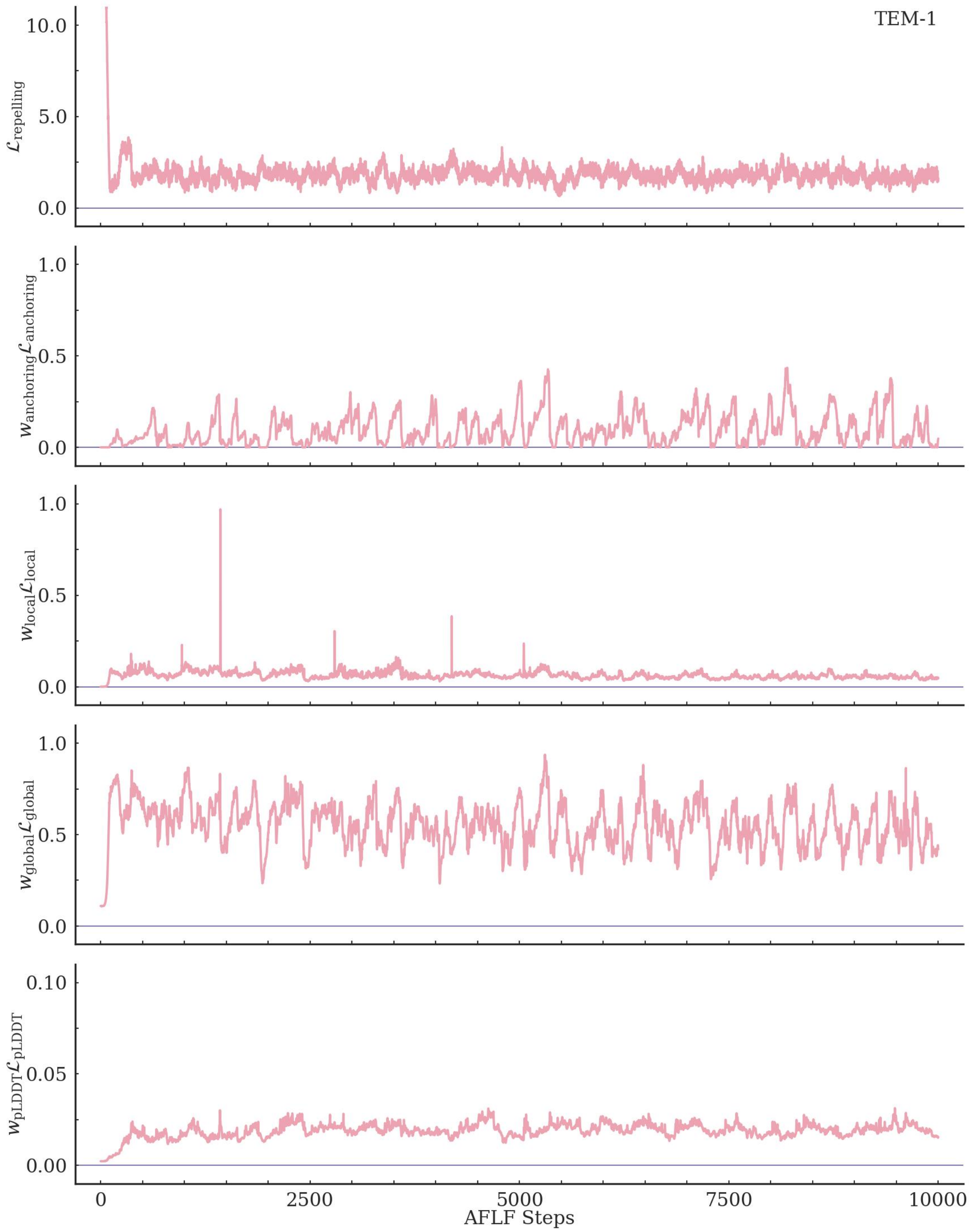

**Supplementary Figure S30.** The evolution of the loss terms along the class A $\beta$-lactamase TEM-1 trajectory.

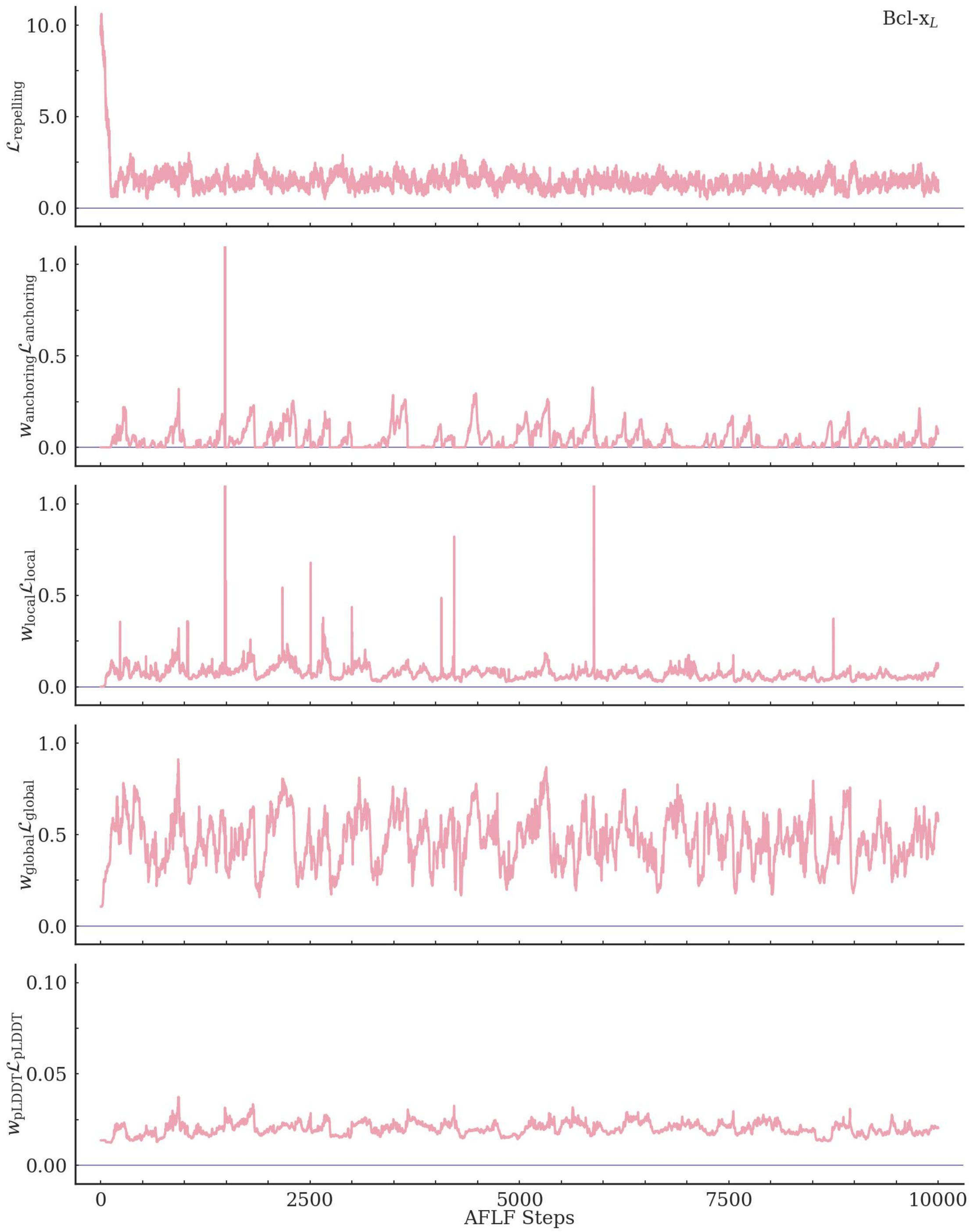

**Supplementary Figure S31.** The evolution of the loss terms along the anti-apoptotic protein Bcl-x$_L$ trajectory.

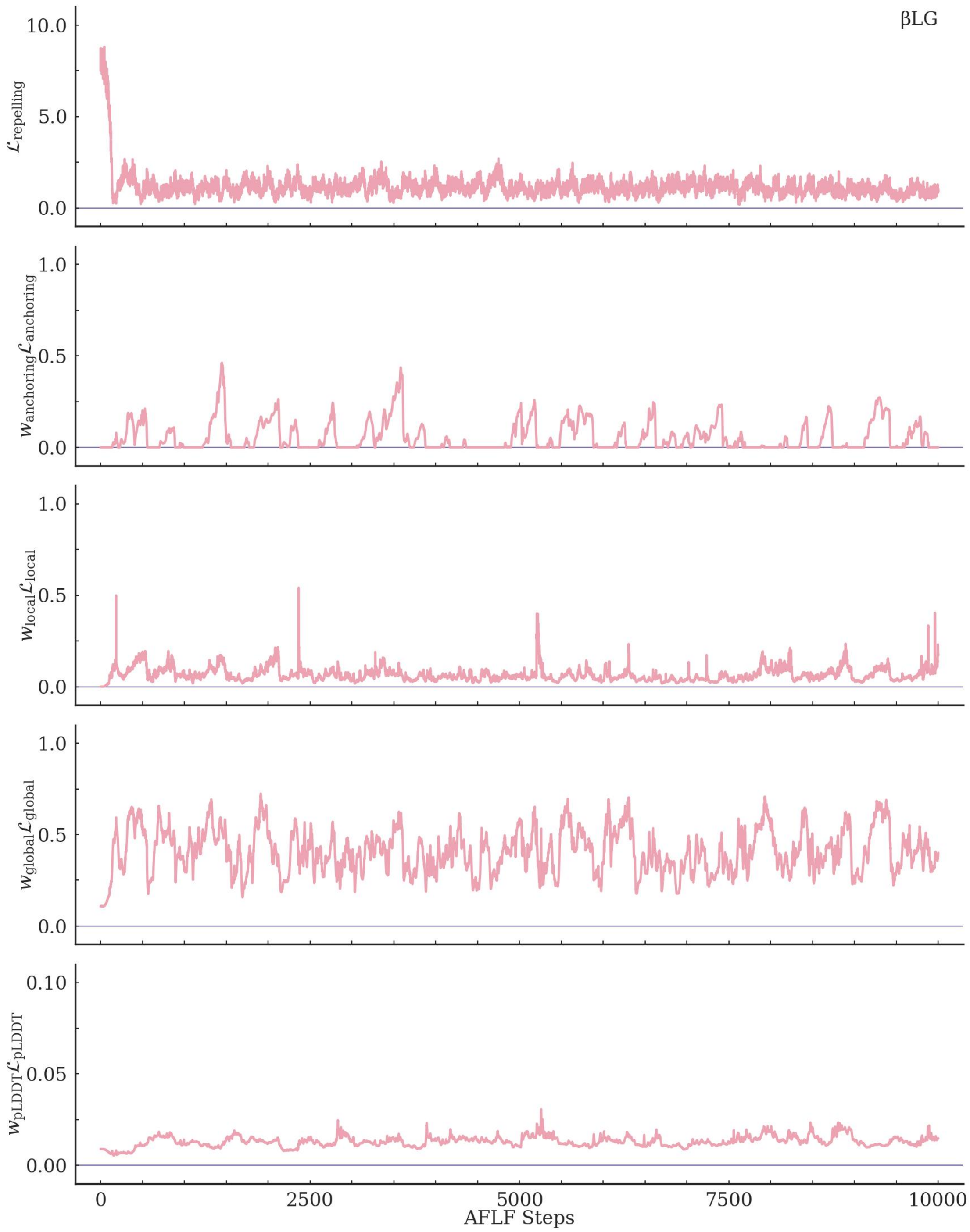

**Supplementary Figure S32.** The evolution of the loss terms along the bovine $\beta$-lactoglobulin ($\beta$LG) trajectory.

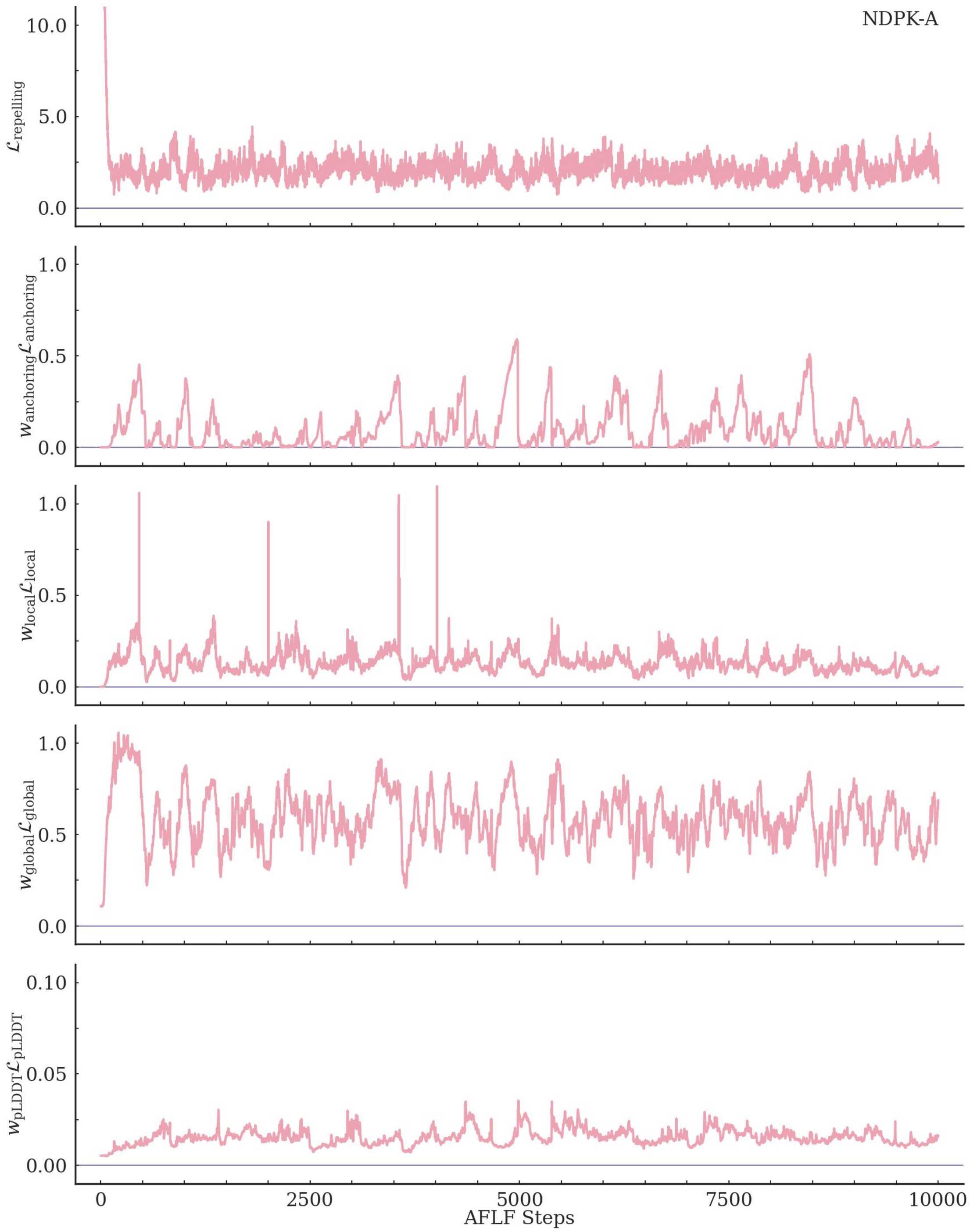

**Supplementary Figure S33.** The evolution of the loss terms along the human nucleoside diphosphate kinase A (NDPK-A) trajectory.

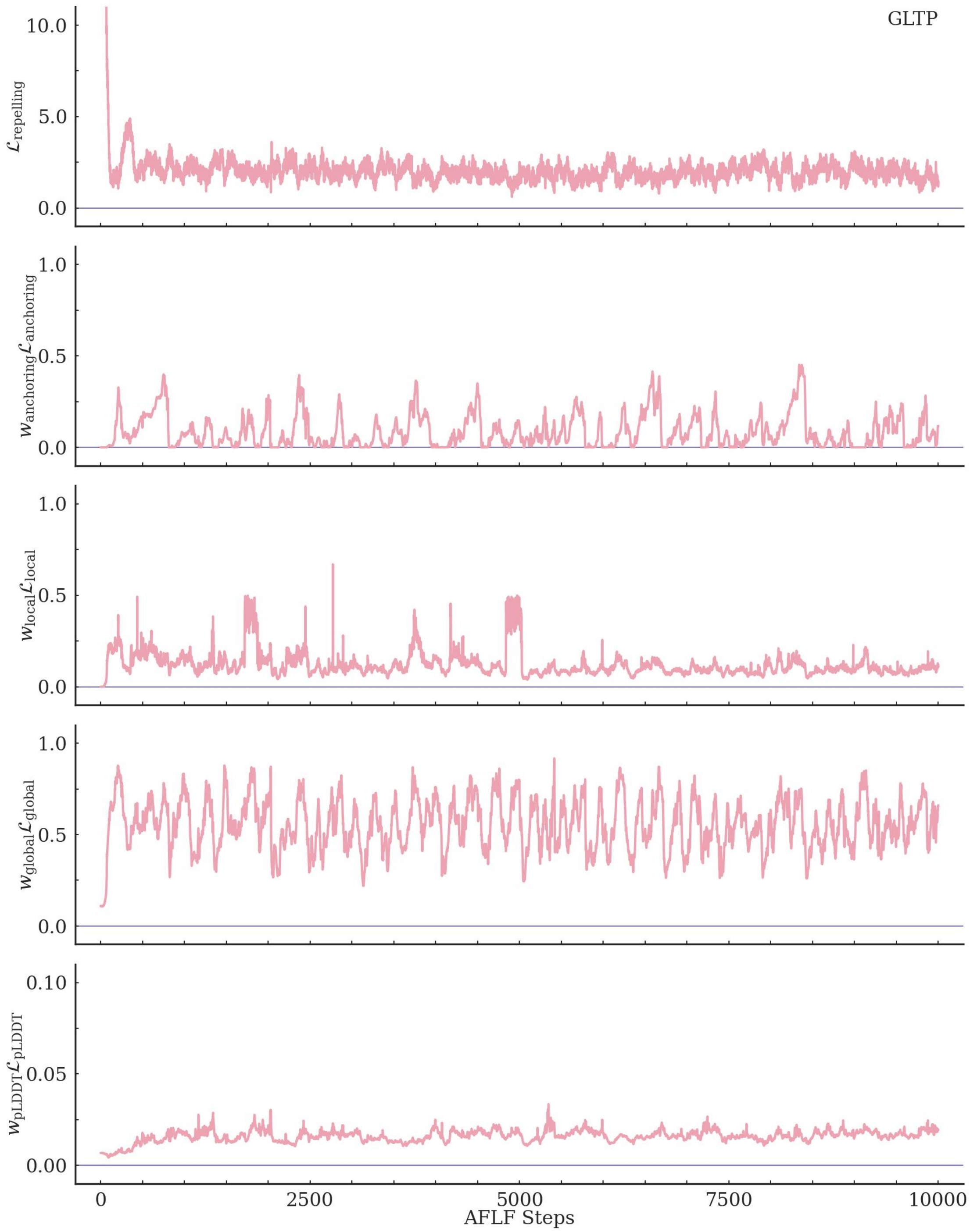

**Supplementary Figure S34.** The evolution of the loss terms along the human glycolipid transfer protein (GLTP) trajectory.

#### 3 Supporting Tables

**Supplementary Table S1.** Experimental and simulated residue-wise B-factors for ubiquitin (unit: $Å^2$).

| Residue | Exp. | MD | AFLF | Residue | Exp. | MD | AFLF |
|---|---|---|---|---|---|---|---|
| MET 1 | 10.38 | 15.32 | 17.17 | ASP 39 | 14.96 | 26.34 | 9.83 |
| GLN 2 | 9.07 | 11.58 | 4.85 | GLN 40 | 10.76 | 21.56 | 7.19 |
| ILE 3 | 5.07 | 6.66 | 3.92 | GLN 41 | 3.87 | 14.06 | 4.10 |
| PHE 4 | 4.68 | 5.00 | 4.35 | ARG 42 | 6.97 | 14.19 | 5.06 |
| VAL 5 | 3.87 | 5.69 | 3.29 | LEU 43 | 3.51 | 8.21 | 2.54 |
| LYS 6 | 6.12 | 8.47 | 4.18 | ILE 44 | 5.55 | 9.14 | 4.20 |
| THR 7 | 7.48 | 28.25 | 5.76 | PHE 45 | 4.70 | 9.44 | 3.78 |
| LEU 8 | 14.15 | 85.86 | 11.23 | ALA 46 | 7.15 | 32.59 | 6.15 |
| THR 9 | 19.24 | 84.06 | 14.41 | GLY 47 | 11.68 | 34.02 | 11.86 |
| GLY 10 | 18.74 | 52.76 | 12.40 | LYS 48 | 8.82 | 27.60 | 14.38 |
| LYS 11 | 11.91 | 23.08 | 12.07 | GLN 49 | 7.18 | 18.31 | 9.68 |
| THR 12 | 9.85 | 12.51 | 5.01 | LEU 50 | 7.41 | 11.93 | 6.22 |
| ILE 13 | 11.84 | 13.33 | 5.40 | GLU 51 | 11.90 | 15.30 | 13.54 |
| THR 14 | 9.63 | 11.06 | 4.55 | ASP 52 | 16.56 | 17.80 | 11.96 |
| LEU 15 | 9.03 | 10.59 | 3.63 | GLY 53 | 11.77 | 19.62 | 11.47 |
| GLU 16 | 11.50 | 12.44 | 3.87 | ARG 54 | 9.05 | 12.87 | 11.13 |
| VAL 17 | 8.85 | 8.60 | 3.95 | THR 55 | 9.03 | 9.92 | 8.16 |
| GLU 18 | 7.08 | 10.28 | 9.98 | LEU 56 | 8.29 | 7.26 | 3.86 |
| PRO 19 | 7.07 | 12.03 | 10.43 | SER 57 | 9.00 | 13.63 | 7.11 |
| SER 20 | 6.28 | 12.20 | 7.98 | ASP 58 | 7.91 | 17.77 | 17.42 |
| ASP 21 | 7.70 | 8.06 | 14.38 | TYR 59 | 8.45 | 13.98 | 9.83 |
| THR 22 | 6.01 | 8.86 | 7.88 | ASN 60 | 13.94 | 14.59 | 11.50 |
| ILE 23 | 9.92 | 7.75 | 5.91 | ILE 61 | 11.78 | 9.39 | 5.69 |
| GLU 24 | 11.81 | 13.49 | 7.39 | GLN 62 | 15.52 | 14.12 | 10.66 |
| ASN 25 | 10.96 | 9.66 | 7.53 | LYS 63 | 11.97 | 11.59 | 13.90 |
| VAL 26 | 5.53 | 6.92 | 4.69 | GLU 64 | 10.94 | 9.77 | 12.72 |
| LYS 27 | 4.14 | 9.86 | 5.84 | SER 65 | 6.90 | 7.28 | 9.23 |
| ALA 28 | 7.74 | 10.05 | 7.49 | THR 66 | 3.80 | 6.22 | 4.39 |
| LYS 29 | 7.90 | 9.63 | 7.53 | LEU 67 | 3.85 | 5.54 | 2.78 |
| ILE 30 | 5.58 | 10.96 | 4.71 | HIS 68 | 4.17 | 7.37 | 2.79 |
| GLN 31 | 8.67 | 13.75 | 6.24 | LEU 69 | 3.97 | 9.25 | 4.06 |
| ASP 32 | 14.01 | 20.86 | 8.09 | VAL 70 | 6.26 | 13.96 | 3.61 |
| LYS 33 | 14.00 | 24.20 | 7.61 | LEU 71 | 16.06 | 24.05 | 4.34 |
| GLU 34 | 10.07 | 24.75 | 8.71 | ARG 72 | 25.83 | 46.43 | 10.43 |
| GLY 35 | 6.29 | 22.87 | 14.10 | LEU 73 | 30.76 | 249.00 | 24.45 |
| ILE 36 | 6.07 | 22.02 | 8.99 | ARG 74 | 35.33 | 267.10 | 39.81 |
| PRO 37 | 9.18 | 26.91 | 10.99 | GLY 75 | 36.07 | 574.13 | 80.29 |
| PRO 38 | 9.08 | 21.09 | 6.39 | GLY 76 | 36.19 | 1212.43 | 323.16 |

**Exp.**: Experimental B-factors (PDB id: 1UBQ);
**MD**: Molecular dynamics simulated B-factors;
**AFLF**: AlphaFold Latent Flooding simulated B-factors.

**Supplementary Table S2.** PDB ids for the protein crystal structures with cryptic binding sites.

| Protein | Abbreviation | Apo state | Holo state |
|---|---|---|---|
| Class A $\beta$-lactamase TEM-1 | TEM-1 | 1JWP | 1PZO |
| Anti-apoptotic protein Bcl-$x_L$ | Bcl-$x_L$ | 3FDL | 2YXJ |
| Bovine $\beta$-lactoglobulin | $\beta$LG | 1BSQ | 1GX8 |
| Human nucleoside diphosphate kinase A | NDPK-A | 3L7U | 2HVD |
| Human glycolipid transfer protein | GLTP | 1SWX | 2EUM |

**Supplementary Table S3.** AFLF parameters used for each task.

| Parameters | Structural Fluctuations | Functional States | Cryptic Cavities | Benchmarks |
|---|---|---|---|---|
| | | $\mathcal{L}_{\text{repelling}}$ | | |
| $\beta_{ij}$ or $\beta_i$ | $\beta_{ij}$ | $\beta_i$ | $\beta_i$ | – |
| $b$ | 1.0 | 0.0 | 10.0 | – |
| $\Delta t$ | 10 | 10 | 10 | – |
| $K$ | 10 | 50 | 40 | – |
| $\varsigma$ | 4.0 | 4.0 | 4.0 | – |
| $\varepsilon_k$ | 0.1 | 0.1 | 0.1 | – |
| $w_k$ | 0.05 | 0.1 | 0.1 | 0.05 |
| $\psi$ dropout rate | 0.5 | 0.5 | 0.5 | – |
| | | $\mathcal{L}_{\text{local}}$ | | |
| $w_{\text{local}}$ | 0.5 | 0.5 | 0.5 | – |
| $\psi$ dropout rate | 0.2 | 0.2 | 0.2 | – |
| | | $\mathcal{L}_{\text{global}}$ | | |
| $\Delta d$ | 2.0 | 2.0 | 2.0 | – |
| $L$ | 10 | 32 | 32 | – |
| $\varsigma'$ | 4.0 | 4.0 | 4.0 | – |
| $w_{\text{global}}$ | 3.0 | 0.5 | 0.5 | – |
| $\psi$ dropout rate | 0.2 | 0.2 | 0.2 | – |
| | | $\mathcal{L}_{\text{anchoring}}$ | | |
| $\delta_{\text{Huber}}$ | 0.1 | 0.1 | 0.1 | – |
| $w_{\text{anchoring}}$ | 1.0 | 1.0 | 3.0 | – |
| $\psi$ dropout rate | 0.0 | 0.0 | 0.0 | – |
| | | $\mathcal{L}_{\text{pLDDT}}$ | | |
| $w_{\text{pLDDT}}$ | 0.1 | 0.1 | 0.1 | – |
| | | AdaM optimizer | | |
| $\eta$ learning rate | 0.002 | 0.002 | 0.003 | – |
| $\beta_1$ | 0.9 | 0.9 | 0.9 | – |
| $\beta_2$ | 0.999 | 0.999 | 0.999 | – |

**Supplementary Table S4.** AFLF computational wall times.[*]

| **Protein** | $N_{res}$ | **JIT[†] (s)** | **Step (s)** | **Protein** | $N_{res}$ | **JIT[†] (s)** | **Step (s)** |
|---|---|---|---|---|---|---|---|
| Representative model systems[§] | | | | | | | |
| UBQ | 76 | 155.48 | 5.57 | AdK | 214 | 179.22 | 16.87 |
| TEM-1 | 263 | 186.36 | 23.26 | Bcl-$x_L$ | 158 | 174.41 | 11.57 |
| $\beta$LG | 162 | 177.08 | 11.77 | NDPK-A | 152 | 161.27 | 10.49 |
| GLTP | 209 | 176.11 | 16.42 | | | | |
| $B$-factor systems[§] | | | | | | | |
| BPTI | 58 | 97.53 | 4.55 | P06654 | 56 | 99.97 | 4.54 |
| P00644 | 149 | 113.41 | 10.75 | P61823 | 124 | 108.03 | 8.59 |
| P00648 | 110 | 111.71 | 7.81 | P62937 | 165 | 104.73 | 12.15 |
| P00698 | 129 | 112.46 | 9.17 | PDZ2 | 96 | 92.72 | 6.62 |
| P01854 | 153 | 115.90 | 11.03 | | | | |
| BioEmu “domain motion” benchmark[¶] | | | | | | | |
| A0A075Q0W3[‡] | 643 | 114.73 | 9.55 | P24182 | 452 | 107.66 | 9.51 |
| A0A0H3AFX3 | 418 | 108.29 | 8.63 | P24182 | 449 | 109.38 | 9.88 |
| A2RJ53 | 590 | 123.77 | 17.05 | P28366[‡] | 844 | 94.55 | 9.79 |
| B0F0C5 | 497 | 115.55 | 12.00 | P28366[‡] | 802 | 119.68 | 10.43 |
| P0205 | 271 | 104.97 | 4.10 | P33284 | 485 | 114.35 | 11.45 |
| P02911 | 238 | 110.85 | 3.31 | P37487 | 309 | 107.42 | 5.08 |
| P03047 | 147 | 105.78 | 1.79 | P67701 | 138 | 106.57 | 1.66 |
| P06766 | 335 | 113.23 | 5.81 | P69441 | 214 | 100.83 | 2.85 |
| P0A8W0 | 263 | 100.89 | 3.92 | P71447 | 221 | 106.24 | 3.00 |
| P0AEQ3 | 226 | 104.88 | 3.07 | Q18A65 | 319 | 117.98 | 5.33 |
| P0DP23 | 148 | 100.61 | 1.76 | Q5F9M1 | 283 | 114.92 | 4.39 |
| P23847 | 507 | 121.06 | 12.39 | Q83VS8 | 351 | 107.98 | 6.34 |
| BioEmu “cryptic pocket” benchmark[¶] | | | | | | | |
| O74933 | 486 | 116.08 | 11.37 | P44542 | 306 | 111.41 | 4.98 |
| P00766 | 245 | 116.11 | 3.52 | P56817 | 411 | 118.97 | 8.37 |
| P02787 | 337 | 130.23 | 5.96 | P60338 | 405 | 117.13 | 8.19 |
| P08037 | 286 | 111.37 | 4.45 | P61586 | 178 | 136.18 | 2.25 |
| P08581 | 312 | 116.84 | 4.79 | P62593 | 263 | 120.03 | 3.92 |
| P08709 | 254 | 118.18 | 3.67 | P79345 | 130 | 113.19 | 1.56 |
| P0A7D4 | 431 | 114.02 | 9.15 | P84080 | 181 | 109.78 | 2.30 |
| P0A7F3 | 153 | 104.33 | 1.87 | P9WPY3 | 184 | 108.41 | 2.23 |
| P0A8V6 | 243 | 109.09 | 3.45 | Q02763 | 327 | 125.84 | 5.58 |
| P0AG16 | 504 | 117.13 | 11.27 | Q16539 | 360 | 114.96 | 6.18 |
| P0DP23 | 148 | 106.75 | 1.77 | Q16658 | 496 | 124.62 | 10.87 |
| P12758 | 253 | 124.74 | 3.67 | Q29495 | 174 | 104.88 | 2.17 |
| P19491 | 263 | 118.88 | 3.92 | Q58L87 | 190 | 114.66 | 2.42 |
| P26281 | 158 | 103.06 | 1.91 | Q8WSF8 | 228 | 113.46 | 3.05 |
| BioEmu “OOD-60” benchmark[¶] | | | | | | | |
| A0A0H3AFX3 | 418 | 108.29 | 8.63 | P50405 | 77 | 104.58 | 0.96 |
| A0A377JKY9 | 92 | 100.45 | 1.14 | P67701 | 138 | 106.57 | 1.66 |
| A0A377JKY9 | 96 | 98.39 | 1.14 | P69801 | 274 | 103.31 | 4.18 |
| B0F0C5 | 497 | 115.55 | 12.00 | P69801 | 266 | 113.12 | 3.99 |
| E1C7U0 | 392 | 105.23 | 7.13 | P69805 | 283 | 114.99 | 4.39 |
| J7QA90 | 88 | 98.41 | 1.05 | Q2FYI5 | 70 | 103.41 | 0.89 |
| K0JNC6 | 215 | 107.26 | 2.87 | Q2FYI5 | 71 | 102.75 | 0.90 |
| P03047 | 147 | 105.78 | 1.79 | Q31PX7 | 125 | 95.45 | 1.49 |
| P0A8W0 | 263 | 100.89 | 3.92 | Q699R5 | 76 | 88.61 | 1.04 |
| P0CL43 | 82 | 93.97 | 1.03 | Q83VS8 | 351 | 107.98 | 6.34 |
| P21926 | 90 | 103.24 | 1.11 | Q9NR30 | 372 | 102.55 | 6.74 |

[*] Wall times measured on a single NVIDIA A40 GPU.
[†] The first AFLF inference step incurs a one-time just-in-time (JIT) compilation overhead.
[‡] To fit within the GPU memory limit, the number of MSA clusters was reduced from the default 508 to 254 for A0A075Q0W3 and P28366 ($N_{res} = 802$), and to 212 for P28366 ($N_{res} = 844$).
[§] AFLF inference from the Evoformer input at layer 1, with LoRA applied to the MSA latent representation.
[¶] AFLF inference from the Evoformer input at layer 40, with LoRA applied to the MSA and pair latent representations.
